# BatR: A novel regulator of antibiotic tolerance in *Pseudomonas aeruginosa* biofilms

**DOI:** 10.1101/2024.04.17.589660

**Authors:** Ainelen Piazza, Harshanie Dasanayaka, Gerhard Saalbach, Carlo Martins, Eleftheria Trampari, Mark A. Webber, Freya Harrison, Jacob G. Malone

## Abstract

*Pseudomonas aeruginosa* is a multidrug-resistant opportunistic human pathogen. Chronic infections are associated with biofilms, conferring resistance to antibiotics and complicating treatment strategies. This study focuses on understanding the role of the uncharacterized gene *PA3049*, upregulated under biofilm conditions. In the context of *P. aeruginosa* biofilms, *PA3049* plays a role in withstanding antimicrobial challenges both *in vitro* and in clinically validated infection models. Under sub-inhibitory concentrations of antibiotic, the deletion of *PA3049* resulted in reduced pyocyanin production and altered abundance of enzymes controlling denitrification, pyoverdine, and hydrogen cyanide biosynthesis. Notably, PA3049 directly interacts with two kinases implicated in stress response, inactivating their active sites. Renamed as the <u>B</u>iofilm <u>a</u>ntibiotic tolerance <u>R</u>egulator (BatR), PA3049 is a key player in *P. aeruginosa* biofilm maintenance and antimicrobial tolerance. These findings contribute to understanding the complex bacterial lifestyle in biofilms, shedding light on a previously uncharacterized gene with significant implications for combating multidrug-resistant infections.

**IMPORTANCE:** *P. aeruginosa* is a multidrug-resistant ESKAPE pathogen that causes chronic biofilm-based infections and is a leading cause of mortality in cystic fibrosis (CF) patients. Understanding the molecular mechanisms underlying *P. aeruginosa* biofilm resilience and antimicrobial resistance is crucial for developing effective therapeutic interventions. This study focuses on characterizing the gene *PA3049*, now known as the *<u>b</u>iofilm <u>a</u>ntibiotic tolerance <u>R</u>egulator* (*batR*). BatR plays a central role within *P. aeruginosa* biofilms, orchestrating adaptive responses to antimicrobial challenges. Our work sheds light on the contribution of *batR* to biofilm biology and its relevance in lung infections, where subinhibitory antibiotic concentrations make BatR pivotal for bacterial survival. By advancing our understanding of *P. aeruginosa* biofilm regulation, this study holds significant promise for the development of innovative approaches against biofilm-associated infections to mitigate the growing threat of antimicrobial resistance.

## INTRODUCTION

*Pseudomonas aeruginosa* is a virulent, opportunistic human pathogen and one of the ESKAPE group of bacteria (1) that are leading causes of multidrug-resistant, nosocomial bacterial infections. *P. aeruginosa* presents a significant challenge in healthcare settings where it causes highly persistent, chronic infections, primarily affecting catheter and cannula implants (2), burn or other major injury victims (3), chronic wounds (4), and immunocompromised patients (5). In 2015, multidrug-resistant *P. aeruginosa* led to an estimated 4,500 attributable deaths and 68,000 infections in the EEA (6), while over 30% of *P. aeruginosa* isolates reported to EARS-Net in 2018 were resistant to at least one regularly monitored group of antimicrobials (7). *P. aeruginosa* is also a major respiratory pathogen, causing both acute and chronic lung infections. Pulmonary infections caused by *P. aeruginosa* are a major cause of mortality and morbidity in people with the genetic disorder cystic fibrosis (8, 9).

Chronic *P. aeruginosa* infections are frequently associated with biofilms, which enable it to evade host immune responses and confer broad resistance/tolerance to antimicrobial agents, complicating treatment strategies. Bacterial biofilms exhibit common traits and phenotypic characteristics, including cell-to-cell communication (quorum sensing), the production and deployment of extracellular polymeric substances and extracellular DNA (eDNA), and the spatially structured control of motility, adhesins and c-di-GMP levels (10).

The upregulation of genes linked to stationary phase adaptation, environmental stress and anaerobiosis further underscores the distinct features associated with biofilm growth (11, 12). Additionally, the spatial arrangement of cells within the biofilm community, exposed to multiple resource gradients, introduces heterogeneity in cell physiology and metabolism that plays an important role in antibiotic tolerance. This diversity includes significant subpopulations of less metabolically active (dormant) cells, which contribute to the notable tolerance of biofilms to antibiotics designed to target active metabolic processes (13). For example, dormant cells within oxygen-depleted zones of *P. aeruginosa* biofilms exhibit lower overall mRNA transcript abundance and increased tolerance to ciprofloxacin and tobramycin (13).

Increased antimicrobial tolerance has also been associated with the global metabolic adaptations that arise in response to the biofilm environment (14). *P. aeruginosa* showcases this adaptability through its ability to grow anaerobically using nitrate as an alternative electron acceptor (15). The relevance of metabolic adaptions in response to antibiotics was underscored by an *in vitro* screen for tobramycin-resistant *P. aeruginosa* PA14 mutants that revealed a significant proportion of resistant mutants with transposon insertions in energy metabolism genes (16).

Although a consensus has emerged on the role of many biofilm-associated traits, the large number of uncharacterized genes reported as differentially expressed under biofilm conditions highlights the substantial gaps that remain in our understanding of this complex bacterial lifestyle (17). Among the minority of upregulated loci in dormant, biofilm-dwelling *P. aeruginosa* cells is the small, uncharacterised gene *PA3049* (13). *PA3049* is predicted to encode a 70-residue cytoplasmic hydrophilin; part of a family of small, extremely hydrophilic, glycine-rich proteins that contribute to desiccation/osmotic stress tolerance in plants and yeast, but whose role in bacteria is less well understood (18).

*PA3049* is annotated as a homolog of the *Escherichia coli* ribosome <u>m</u>odulator factor (RMF) (13). RMF, a ribosomally associated protein, facilitates ribosome hibernation by associating with 100S ribosome dimers and modulates *E. coli* translation during the stationary phase (19). Unlike other ribosome components or associated proteins, *rmf* expression is inversely dependent on growth rate, indicating its potential involvement in the bacterial stress response (19). Classified as a hydrophilin, RMF accumulates under hyperosmotic stress conditions (18). RMF deletion in *E. coli* is linked to decreased aminoglycoside tolerance, potentially due to the location of its ribosome binding site (20). However, while RMF is crucial for ribosome hibernation in *E. coli* (21), PA3049 apparently does not fulfil a similar function in *P. aeruginosa* (22). Deleting *PA3049* in *P. aeruginosa* compromises membrane integrity in biofilm-forming cells, suggesting a potential role in maintaining cell viability for the dormant subpopulation (13). Nonetheless, despite its conservation in sequenced *P. aeruginosa* strains and likely functionality in biofilms, the precise role of *PA3049* in *P. aeruginosa* biofilm formation remains unknown (22).

Here, we used a combination of bioinformatics, infection biology and molecular microbiology to determine the distribution of *PA3049* across bacterial genomes, its importance for host infection and biofilm formation, its contribution to antimicrobial tolerance and its mechanism of action in *P. aeruginosa*. *PA3049*-like genes are widespread in γ-proteobacteria, with a substantial degree of sequence and structural divergence predicted between the Pseudomonadales and Enterobacterales.

*PA3049* plays an important role in the formation of biofilm architecture and enabling established biofilms to withstand antimicrobial challenge. These global traits were traced to a narrow set of phenotypic and protein abundance shifts, with *PA3049* deletion leading to reduced pyocyanin production and altered abundance of enzymes controlling denitrification, pyoverdine and hydrogen cyanide biosynthesis. Finally, we showed that PA3049 interacts directly with two different kinases; SrkA and GlpK that have recently been linked to antibiotic tolerance, suggesting an *in vivo* mechanism of action. Given the importance of *PA3049* for *P. aeruginosa* antimicrobial tolerance and biofilm maintenance, we renamed this gene <u>B</u>iofilm <u>a</u>ntibiotic tolerance <u>R</u>egulator (BatR).

## RESULTS

### *PA3049* homologs are widespread in γ-proteobacteria

To uncover the role of *PA3049* (*batR*) in *P. aeruginosa*, we first compared its predicted structure with that of *E. coli* RMF and assessed the wider distribution of *batR*-like genes in bacterial genomes. An alignment of BatR and RMF sequences, sharing 49.09% identity, shows a 15-residue C-terminal extension on BatR that is absent from RMF (Fig. 1a). AlphaFold three-dimensional protein structure prediction of BatR highlighted a marked difference in the fold of the protein C-terminus compared to its *E. coli* homolog. BatR contains a predicted α-helix of 12 residues at the C terminus (red boxed) in addition to the two α helices connected by a 13-amino acid linker region that comprise *E. coli* RMF (Fig. 1b).

**FIG 1.**
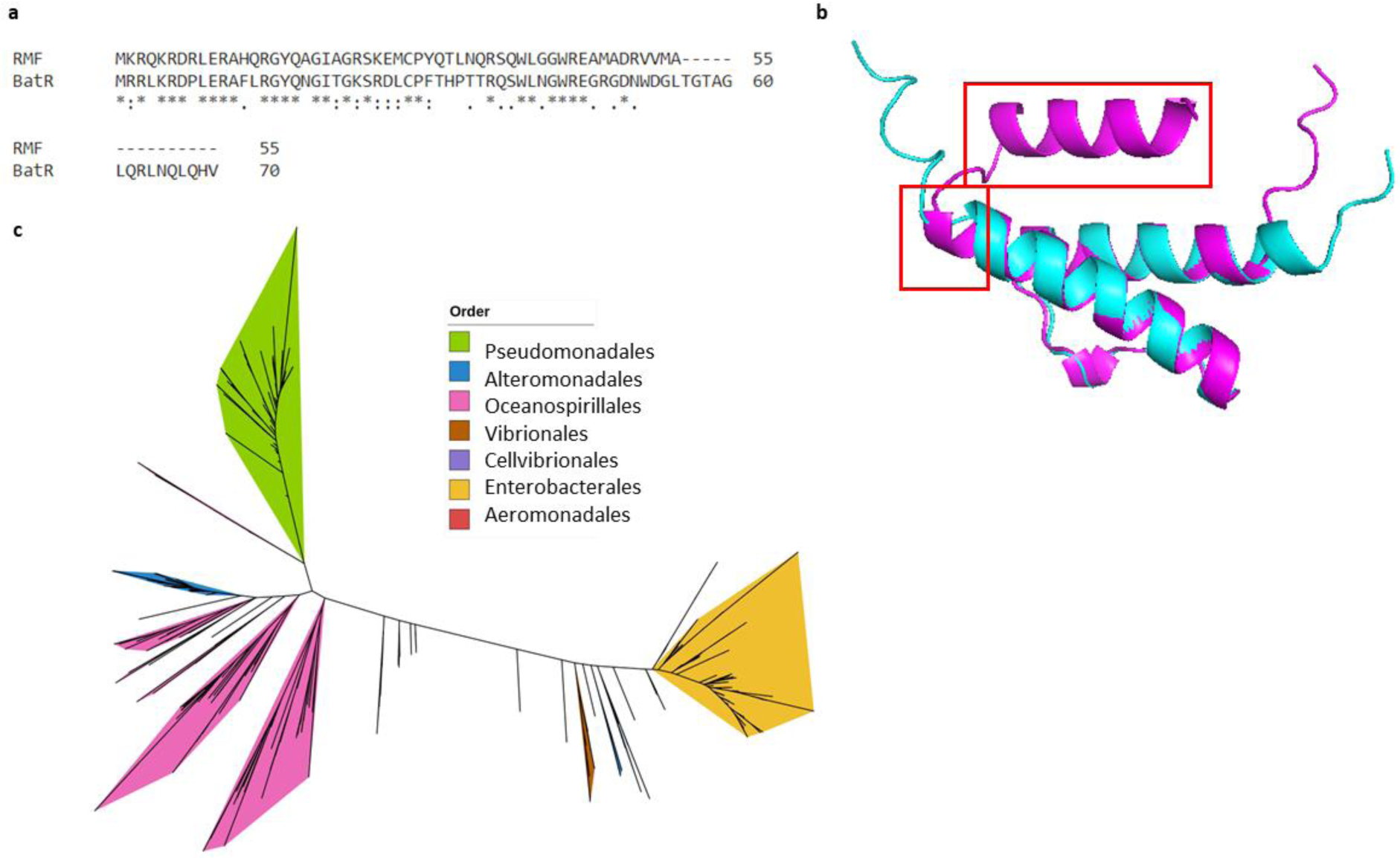
a. Sequence alignment of BatR and RMF. The amino acid sequences were aligned using ClustalW and black asterisks mark conserved residues in both proteins. **b. AlphaFold model of BatR** (magenta), overlaid onto the structure of *E. coli* RMF (cyan). Note the additional alpha-helix predicted for BatR (red boxed) at the C-terminus of the predicted structure. **c. The phylogenetic relationship between BatR/RMF homologs.** The tree is based on 765 proteins homologous to **BatR/**RMF from γ-proteobacteria.

A phylogenetic tree of RMF homologs revealed that they are confined to the γ-proteobacteria class (Fig. 1c), with genes associated with RMF showing distinct evolutionary paths across diverse bacterial orders. Notably, *batR* in Pseudomonadales forms a distinct cluster that diverges significantly from other bacterial orders. This divergence is particularly evident for the Enterobacterales, including *E. coli* and *Yersinia pestis*; and Vibrionales including *Vibrio cholerae,* where RMF homologs have been shown to play roles in ribosome hibernation (23). While *rmf* and *batR* appear to share an ancestral root, their divergent structure and phylogeny, supported by the existing literature (22) are consistent with an alternative functional role for BatR.

### *batR* protects *P. aeruginosa* biofilms from sub-inhibitory concentrations (SIC) of antibiotics *in vitro*

Given the heightened abundance of *batR* transcripts in dormant *P. aeruginosa* cells within biofilms (13), we assessed the importance of *batR* in biofilm formation and its effects on antimicrobial tolerance. To do this, we generated a non-polar deletion mutant (Δ*batR*) in PAO1 and examined the response of the mutant to antimicrobial agents when grown in liquid and on solid agar. Assays were conducted using several different antibiotic classes, targeting distinct metabolic processes: aminoglycosides (gentamicin, GENT), β-lactams (piperacillin, PIP) and quinolones (ciprofloxacin, CIP).

Consistent with previous findings (22), the deletion of *batR* did not impact bacterial viability or result in growth impairment in shaking cultures exposed to SIC of tested antimicrobials (see Fig. S1). Additionally, the minimal inhibitory concentration (24) of all three tested antibiotics was unaffected in Δ*batR* (Table 1). Interestingly however, Δ*batR* showed a small but significant increase in sensitivity to PIP and CIP (12-17%), when grown on a solid surface (Fig 2a).

**FIG 2.**
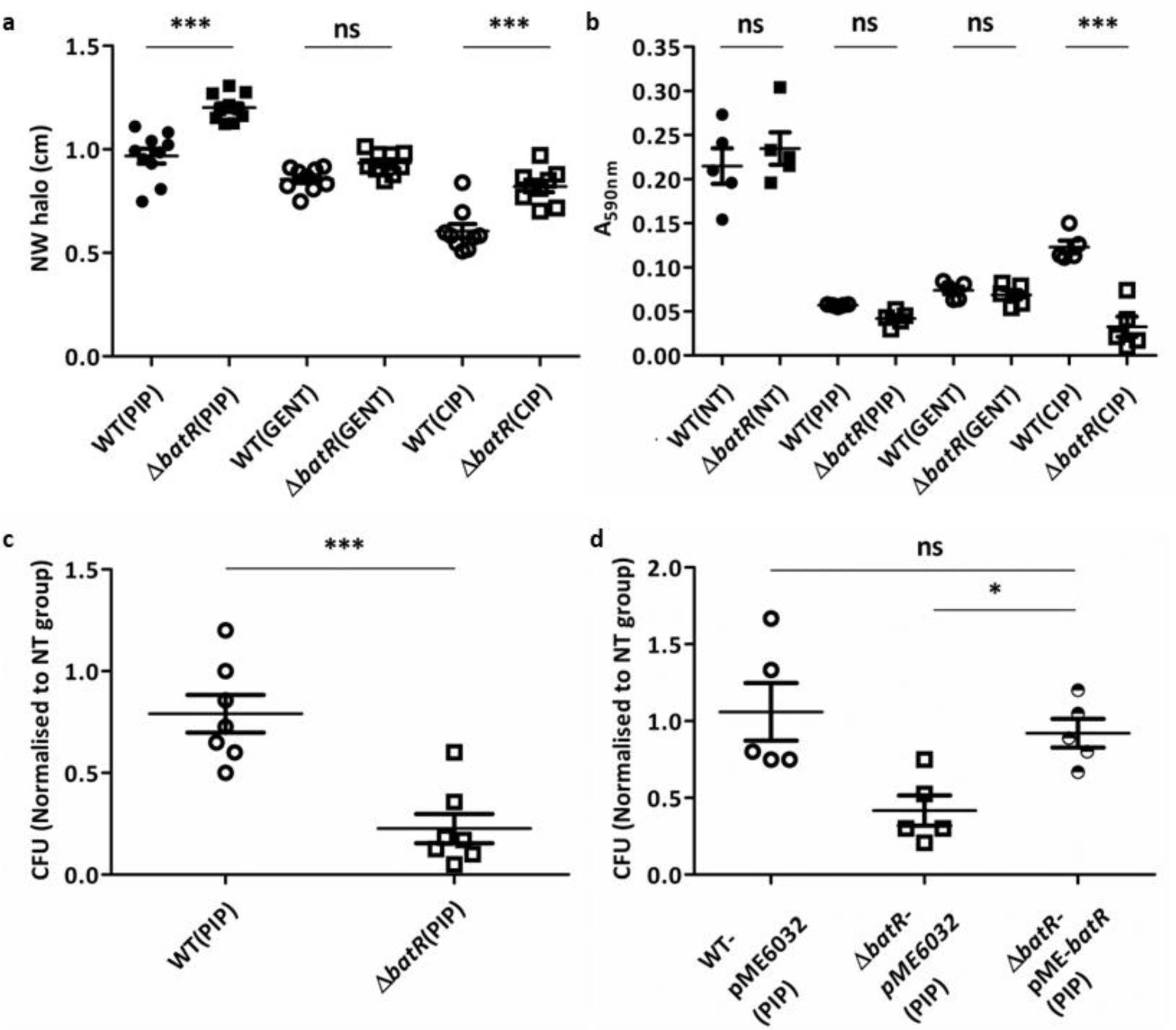
a. Antibiotic disk diffusion assay. Results are represented as the normalized width of the antimicrobial halo (NW halo), calculated as described in (55). Differences in inhibition were significant between WT and Δ*batR* strains (F_5,50_=52.52, P<0.0001) and post hoc analysis showed significant differences between WT and Δ*batR* strain under PIP and CIP treatments (p<0.0001 ***) and non-significant (ns) effect with GENT. **b. Biofilm formation.** Crystal Violet absorbance (A_590nm_) after 24-h static growth in LB medium. Treated samples were incubated in the presence of SIC of antibiotics PIP, GENT, and CIP. NT=not treated. ANOVA F_7,32_=49.35; p<0.0001, post hoc analysis showed this was only significant for CIP treatment (p<0.0001 ***) **c. Glass Beads Biofilm survival**. Bacterial recovery (CFU/bead) from established biofilms grown on glass beads for 24h following treatment with SIC of PIP for 90 min. **d**. **Glass Beads Biofilm survival complementation**. Strains: WT PAO1 strain carrying the empty vector pME6032 (WT-pME6032) and Δ*batR* either carrying the empty vector pME6032 (Δ*batR*-pME6032) or overexpressing *batR* (Δ*batR-*pME-*batR*). ANOVA F_2,12_=6.414; p=0.0127, post hoc analysis showed this was significant for WT-pME6032 (PIP) vs Δ*batR*-pME6032 (PIP) and for Δ*batR*-pME6032 (PIP) vs Δ*batR*-pME-*batR* (PIP) (p<0.05*).

**Table 1.** Minimal inhibitory concentration (MIC)

| Antimicrobial | MIC<sup>a</sup> |  |
| --- | --- | --- |
| | WT PAO1 | $\Delta batR$ PAO1 |
| Piperacillin | 64 | 64 |
| Gentamycin | 8 | 8 |
| Ciprofloxacin | 0.5 | 0.5 |

<sup>a</sup> Expressed in µg/mL unless indicated otherwise.

The comparison of PAO1 WT and Δ*batR* cultures grown in liquid medium (24) under static conditions revealed no measurable effect on biofilm formation (Fig. 2b). However, the addition of SIC of CIP led to a significant reduction in biofilm formation for the Δ*batR* strain compared to WT PAO1 (Fig. 2b). Given the initial observation that sensitivity to both PIP and CIP was affected on solid surfaces (Fig. 2a), we used an alternative, glass bead biofilm model (25), to assess the survival of cells within established biofilms challenged with PIP. This model revealed a substantial reduction in the survival of established Δ*batR* biofilms compared to WT PAO1, for samples exposed to SIC of PIP (Fig. 2c). Notably, the Δ*batR* phenotype could be fully rescued by the heterologous expression of *batR* (Fig. 2d). Considering the different mode of action of the antibiotics tested in this study, our results suggest that the contribution of BatR to *P. aeruginosa* antimicrobial tolerance may occur through a general, rather than a drug-specific mechanism. A simple explanation for these phenotypes could be a nonspecific change in membrane permeability. However, we discarded this hypothesis after quantifying the intracellular concentration of resazurin, a fluorescent dye used as a permeability and efflux marker (Fig. S2), where little difference was seen in the response of WT and Δ*batR* strains.

### BatR affects the ability of *P. aeruginosa* to withstand antimicrobial challenge in established biofilms and clinically validated infection models

To assess the clinical significance of BatR in *P. aeruginosa*, we employed the established *ex vivo* pig lung (EVPL) model (26) to simulate *P. aeruginosa* biofilm infections in CF bronchioles. This model comprises two environments: the bronchiolar lung tissue surface and the SCFM (Synthetic Cystic Fibrosis Medium) to mimic the luminal mucus in the human CF lung (27). We first determined the efficiencies of the antibiotics PIP, CIP, and GENT to clear EVPL infections. Under our test conditions, PAO1 biofilms were highly tolerant to PIP treatment (Fig. S3), precluding its use in these assays. Therefore, we assessed survival of the biofilms challenged with CIP at a SIC of 16 µg/ml. We recovered the biofilms at 2 d and 7 d postinfection (PI) to determine CFU per EVPL section. Our results show that BatR not only contributes to CIP tolerance, but also plays a key role in biofilm establishment within the EVPL model at 2 d PI (Fig. 3a). However, by 7 d PI, while *P. aeruginosa* remained viable on the lung tissue, *batR* did not appear to contribute significantly to biofilm formation or CIP tolerance, although there was a decrease in tolerance to CIP treatment in the WT strain (Fig. 3a). These findings suggest a role for BatR in biofilm establishment and antibiotic tolerance in *P. aeruginosa* infections, particularly in early stages of CF. We further explored these phenotypes in an *in vitro* biofilm model designed to mimic the conditions found in chronic wounds (28). Survival comparisons in biofilms without CIP challenge revealed no differences between WT and Δ*batR* strains. However, significant differences were observed for PAO1 Δ*batR* biofilms treated with different concentrations of CIP, compared to WT PAO1 in this model (Fig. 3b). Next, we examined the biofilm architecture in infected lung pieces, with and without SIC of CIP in the EVPL + SCFM. Replica sets of infected lung pieces were fixed at both 2 d and 7 d PI and sections stained with H & E (Haematoxylin & Eosin) to visualise the total biofilm mass and tissue architecture. At 7 d PI, the EVPL biofilms show the typical “sponge”-like appearance consisting of extracellular matrix punctuated by gaps, resembling CF biofilms observed *in vivo* (29–31). However, there were clear qualitative differences in the biofilm observed between the Δ*batR* and WT strains, as shown in Fig. 4. The matrix covering the biofilm structure appeared thicker in the WT biofilm when challenged with SIC of CIP, something that is not observed in the biofilm formed by the Δ*batR* strain (Fig. 4). These observations suggest that *batR* may play a role in influencing biofilm architecture, particularly in response to SIC of antibiotics.

**FIG 3.**
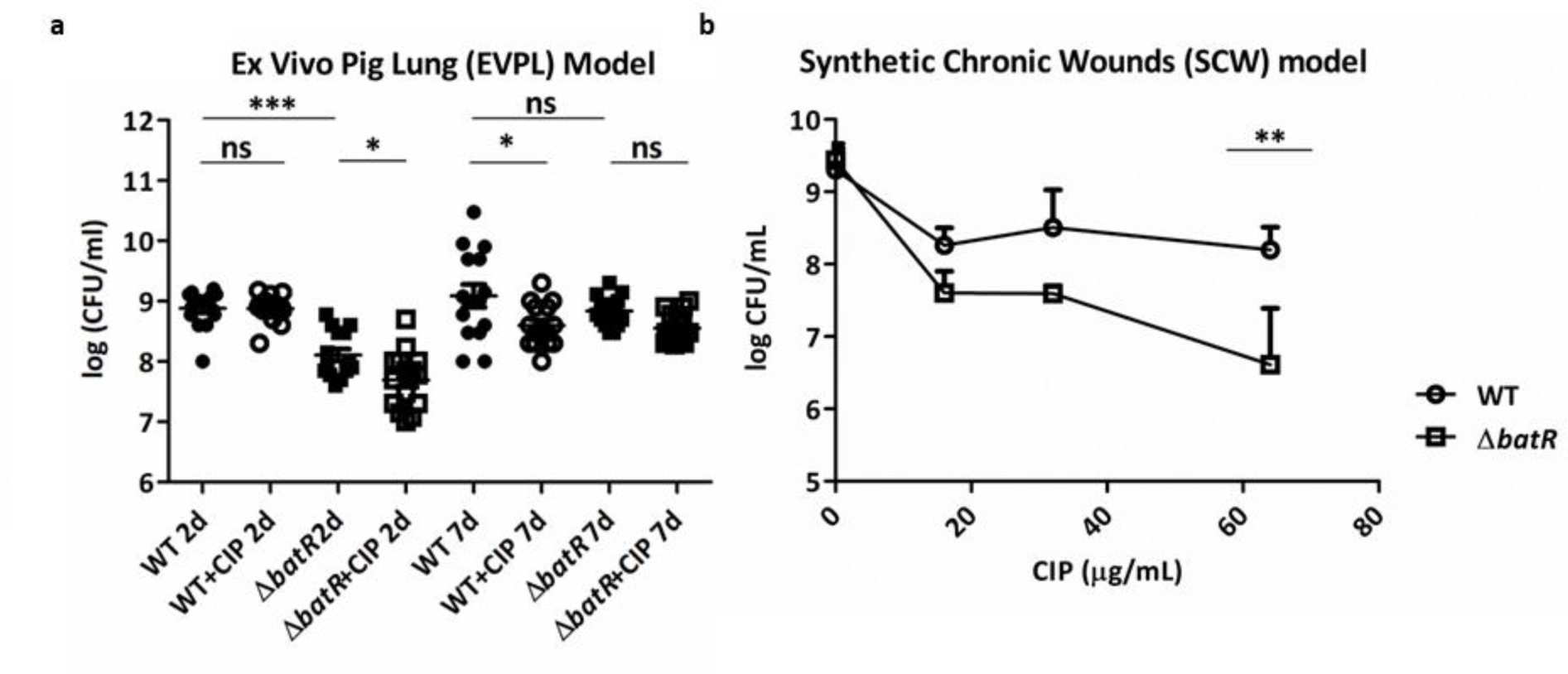
**a. *Ex Vivo* Pig Lung (EVPL) Model**. Growth of *P. aeruginosa* PAO1 strain (WT) and Δ*batR*, on 15 pieces of EVPL bronchiole (five replicate pieces of tissue infected per strain from five independent lungs) plus Synthetic Cystic Fibrosis Medium (SCFM), with and without 16 µg/ml CIP treatment. Colony forming units (CFU) were retrieved from biofilms after 2 d and 7 d growth at 37°C. Bars denote mean for each genotype across all five lungs, and asterisks denote a significant difference under that condition. ANOVA F_7,112_=19.90; p<0.0001, post hoc analysis showed significant differences for WT 2d vs Δ*batR* 2d; WT 2d vs Δ*batR* +CIP 2d; Δ*batR* 2d vs WT+CIP 2d, (p<0.0001 ***); and for WT 7d vs WT+CIP 7d; WT 7d vs DbatR+CIP 7d (p<0.05 *). **b. Synthetic Chronic Wound (SCW) Model**. Viability of *P. aeruginosa* PAO1 (WT) and the Δ*batR* strain living in established biofilms (24 h), treated with different concentrations of CIP in the cell suspension on top of the matrices. Dots represent the average from three biological replicates and error bars indicate the standard deviation. ANOVA found a significant difference in biofilm survival between WT and Δ*batR* under 64 µg/ml CIP treatment.

**FIG 4.**
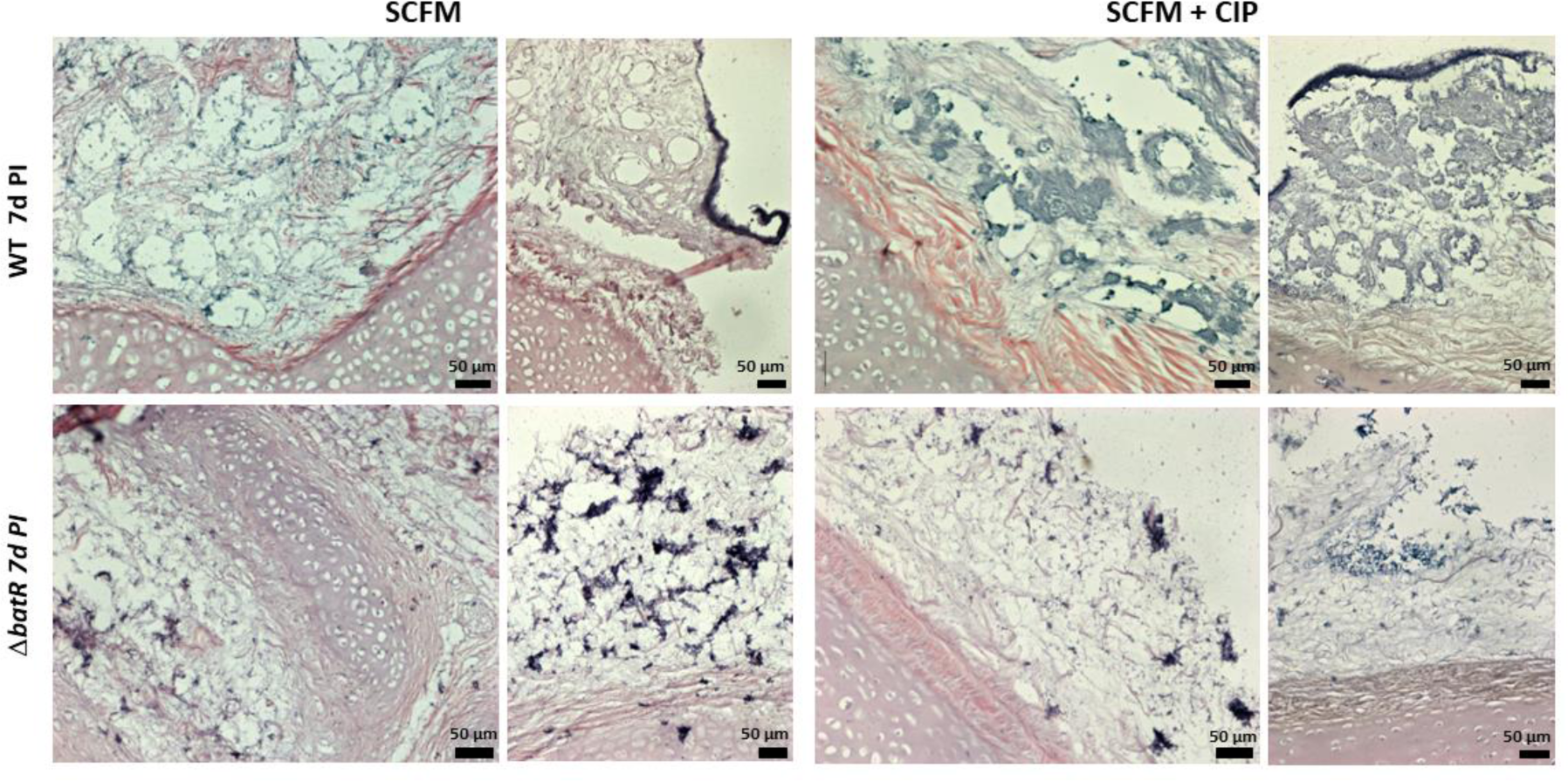
Haematoxylin and eosin (H & E) stained sections of EVPL bronchiolar tissue with SCFM medium infected with *P. aeruginosa* at 7 d post infection. EVPL was infected with *P. aeruginosa* PAO1 WT and Δ*batR*, with uninfected tissue as a negative control. The x20 magnification images from the sections are shown here for non-treated tissues (SCFM) and treated with CIP (SCFM + CIP). The cartilage and tissue surface (lower half of each image) stain pink and the bacterial biofilm stain purple, including the bacterial cells and biofilm matrix. Representative images of phenotypes at day 7 PI are shown here, but the same results were observed for all biological replicates analysed.

### BatR induces the production of specific virulence factors in the EVPL model under CIP challenge

To understand the impact of BatR on *P. aeruginosa* biology in the EVPL model, we quantified the production of key virulence-associated exoproducts (27). After washing the lung tissues twice in PBS to remove planktonic cells, we measured virulence factor production by biofilms growing on the lung tissue surface. Despite non-significant changes in siderophore abundance (pyoverdine and pyochelin, Fig. S4), a notable increase in blue colouration, indicative of pyocyanin production, was evident in the WT strain compared to Δ*batR* when treated with CIP on day 2 (Fig. 5 a,b).

**FIG 5.**
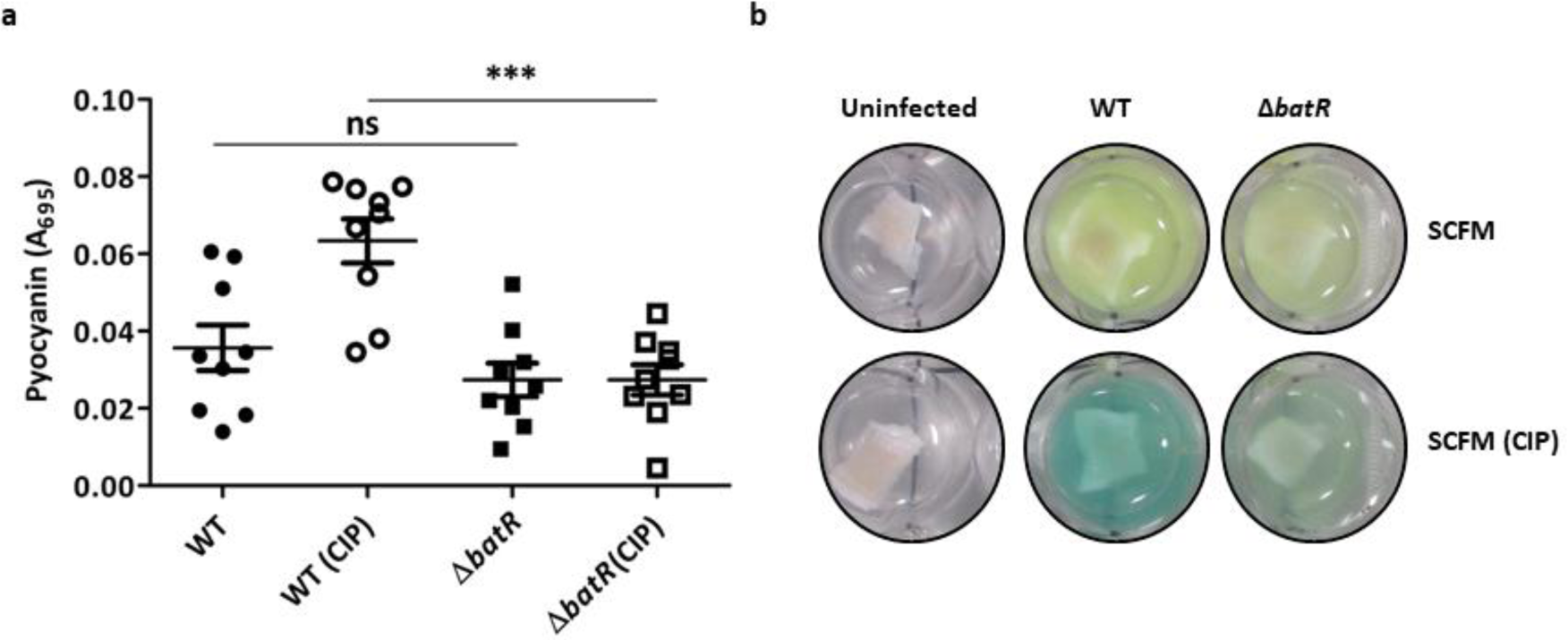
**a. Production of pyocyanin (A_695_) by *P. aeruginosa* WT and Δ*batR* in the EVPL model**. Significant differences in pyocyanin production between WT and Δ*batR* under CIP treatment (ANOVA F_3,32_ = 16.10, p<0.0001***), post hoc analysis showed significant differences (p<0.0001***) for WT vs WT (CIP), WT vs Δ*batR* (CIP); Δ*batR* vs WT (CIP) and Δ*batR* vs Δ*batR* (CIP). **b. *P. aeruginosa* biofilms on squares of bronchiolar tissue.** The blue pigmentation is typical of *P. aeruginosa* and is a mixture of the exoproducts pyoverdine and pyocyanin; note the increased intensity of blue colour in the WT strain grown in SCFM exposed to CIP.

### BatR induces specific changes in the PAO1 proteome under antimicrobial challenge

Next, to investigate the physiological changes associated with BatR during antibiotic challenge in *P. aeruginosa* biofilms, we conducted a comparative proteomic analysis between the WT and Δ*batR* PAO1 strains, in the presence and absence of SIC of the antibiotic PIP. High-resolution mass spectrometry following TMT labelling detected an average of 4,581 individual proteins in each sample (S1 Data), representing ∼80% of the predicted total *P. aeruginosa* PAO1 proteome (32). BatR was exclusively detected in the WT PAO1 strain. Surprisingly, limited differences were detected between the proteomes of the two strains (Fig. S5, S1 Data), suggesting that the impact of *batR* deletion is quite specific under the conditions tested.

Of 21 proteins decreased in the Δ*batR* strain in the absence of PIP challenge (Table 2), seven (PA0617-PA0633) are components of a predicted bacteriophage. In addition, we saw reduced abundance of proteins involved in transport & metabolism; two heat shock proteins and four proteins of unknown function in the Δ*batR* strain. Conversely, 14 proteins were increased in the Δ*batR* strain, including Rubredoxin-1, Type VI secretion system components and the Phenazine-1-carboxylate N-methyltransferase PhzM (Table 3).

**Table 2.** COG pathway analysis of proteins decreased in the Δ*batR* strain.

| Locus tag | Gene | Product | Cog |
| --- | --- | --- | --- |
| **General function prediction only** |  |  |  |
| PA0617 |  | Probable bacteriophage protein | General function prediction only |
| PA0618 |  | Probable bacteriophage protein | General function prediction only |
| PA0619 |  | Probable bacteriophage protein | General function prediction only |
| PA0620 |  | Probable bacteriophage protein | General function prediction only |
| PA0623 |  | Probable bacteriophage protein | General function prediction only |
| PA0626 |  | Phage tail protein | General function prediction only |
| PA0633 |  | Phage tail protein | General function prediction only |
| PA0803 |  | VOC domain-containing protein | Function unknown |
| PA2853 |  | Major outer membrane lipoprotein (Outer membrane lipoprotein I) | Function unknown |
| PA3796 |  | DUF937 domain-containing protein | Function unknown |
| PA1625 |  | AI-2E family transporter | General function prediction only |
| **Transport & Metabolism** |  |  |  |
| PA3224 |  | 3-phosphoglycerate kinase | metabolism |
| PA3280 | *oprO* | Porin O | Inorganic ion transport and metabolism |
| PA3113 | *trpF* | N-(5'-phosphoribosyl) anthranilate isomerase (PRAI) | Amino acid transport and metabolism |
| PA1906 |  | HIT family protein | Nucleotide transport and metabolism<br>Carbohydrate transport and metabolism |
| PA2594 |  | PBPb domain-containing protein | Inorganic ion transport and metabolism |
| PA2941 |  | VWFA domain-containing protein | Coenzyme metabolism |
| **Translation, Posttranslational modification, protein turnover, chaperones** |  |  |  |
| *PA3049* | *rmf/batR* | **Ribosome modulation factor** | **Translation, ribosomal structure, and biogenesis** |
| PA5195 | *yrfH* | Heat shock protein 15 | Translation, ribosomal structure, and biogenesis |
| PA4762 | *grpE* | Protein GrpE (HSP-70 cofactor) | Posttranslational modification, protein turnover, chaperones |
| **Membrane** |  |  |  |
| PA1041 |  | Probable outer membrane protein | Cell envelope biogenesis, outer membrane –<br>Cell motility and secretion |

**Table 3.** COG pathway analysis of proteins increased in the Δ*batR* strain.

| Locus tag | Gene | Product | Cog |
| --- | --- | --- | --- |
| **Energy production and conversion** |  |  |  |
| PA5351 | *rubA1* | Rubredoxin-1 (Rdxs) | Energy production and conversion |
| **Lipid Metabolism** |  |  |  |
| PA0286 | *desA* | Delta-9 fatty acid desaturase | Lipid metabolism |
| PA3157 | *wbpC* | Probable acetyltransferase | Lipid metabolism |
| **Type VI secretion system** |  |  |  |
| PA1658 | *hsiC2* | Type VI secretion system contractile sheath large subunit | Intracellular trafficking, secretion, and vesicular transport |
| PA2366 | *hsiC3* | Uricase PuuD | Intracellular trafficking, secretion, and vesicular transport |
| PA2367 | *hcp* | Type VI secretion system tube protein | Intracellular trafficking, secretion, and vesicular transport |
| PA0071 | *tagR1* | FGE-sulfatase domain-containing protein | Hcp secretion island I (HSI-I) type VI secretion system |
| PA2361 | *icmF3* | IcmF-related_N domain-containing protein | Intracellular trafficking, secretion, and vesicular transport |
| **Secondary metabolites biosynthesis, transport, and catabolism** |  |  |  |
| PA4209 | *phzM* | Phenazine-1-carboxylate N-methyltransferase | phenazine biosynthetic process |
| PA4143 | *cyaB* | Probable toxin transporter | transmembrane transport |
| **Inorganic ion transport and metabolism** |  |  |  |
| PA1436 |  | Probable Resistance-Nodulation-Cell Division (RND) efflux transporter |  |
| **Signal transduction mechanisms** |  |  |  |
| PA4154 | *ygiM* | SH3b domain-containing protein | Signal transduction mechanisms |
| **Unknown function** |  |  |  |
| PA4297 | *tadG* | TadG | Function unknown |
| PA2030 |  | Transmembrane protein | Function unknown |

For cells subjected to PIP challenge (S1 Data), *batR* deletion is associated with significantly decreased abundance of 11 proteins (Table 4), including components of the Hydrogen cyanide (HCN) and Pyoverdine (PVD) synthesis pathways, quorum sensing and transcriptional regulators, and iron transport proteins. Deletion of *batR* significantly increased abundance of only twelve proteins under PIP challenge. These proteins play roles in energy production and conversion, especially nitrate respiration, primary metabolism, and Type VI secretion (Table 5).

**Table 4.** COG pathway analysis of proteins decreased in the Δ*batR* strain under PIP challenge.

| Locus tag | Gene | Product | Cog |
| --- | --- | --- | --- |
| **Quorum sensing** |  |  |  |
| PA0998 | *pqsC* | 2-heptyl-4(1H)-quinolone synthase subunit | acyl-carrier-protein - Secondary metabolites biosynthesis, transport, and catabolism - Lipid metabolism |
| PA4341 |  | Probable transcriptional regulator | Transcription |
| **HCN production** |  |  |  |
| PA2193 | *hcnA* | Hydrogen cyanide synthase subunit HcnA | Energy production and conversion - General function prediction only |
| PA2194 | *hcnB* | Hydrogen cyanide synthase subunit HcnB | General function prediction only |
| **pyoverdine synthesis** |  |  |  |
| PA2386 | *pvdA* | L-ornithine N(5)-monooxygenase | Secondary metabolites biosynthesis, transport, and catabolism |
| PA2402 | *pvdI* | Pyoverdine peptide synthetase | Secondary metabolites biosynthesis, transport, and catabolism - Lipid metabolism - Secondary metabolites biosynthesis, transport, and catabolism |
| PA2413 | *pvdH* | L-2,4-diaminobutyrate:2-ketoglutarate 4-aminotransferase | Amino acid transport and metabolism<br>- Coenzyme metabolism |
| **Transcription/Translation** |  |  |  |
| PA2511 | *antR* | Probable transcriptional regulator | Transcription |
| **PA3049** | ***rmf/batR*** | ***Ribosome modulation factor*** | ***Translation, ribosomal structure, and biogenesis*** |
| **Transport** |  |  |  |
| PA2512 | *antA* | Anthranilate dioxygenase large subunit | Inorganic ion transport and metabolism |
| PA4358 | *feoB* | Ferrous iron transport protein B | Inorganic ion transport and metabolism |

**Table 5.** COG pathway analysis of proteins increased in the Δ*batR* strain under PIP challenge.

| Locus tag | Gene | Product | Cog |
| --- | --- | --- | --- |
| **Type VI Secretion System Proteins** |  |  |  |
| PA0263;<br>PA1512;<br>PA5267 | *Hcp* | Major exported protein | Intracellular trafficking, secretion, and vesicular transport |
| PA2367 | *hcp3* | Type VI secretion system tube protein Hcp | Intracellular trafficking, secretion, and vesicular transport |
| **Metabolism** |  |  |  |
| PA1897 |  | Fatty acid hydroxylase domain-containing protein | Lipid metabolism |
| **Energy production - Nitrate Respiration Proteins** |  |  |  |
| PA0520 | *nirQ* | Denitrification regulatory protein | General function prediction only |
| PA0524 | *norB* | Nitric oxide reductase subunit B | Posttranslational modification, protein turnover, chaperones - Inorganic ion transport and metabolism |
| PA3391 | *nosR* | Regulatory protein | Transcription - Energy production and conversion |
| PA3392 | *nosZ* | Nitrous-oxide reductase precursor | Energy production and conversion |
| **Energy production** |  |  |  |
| PA5351 | *rubA1* | Rubredoxin-1 | Energy production and conversion |
| **Transport** |  |  |  |
| PA1436 |  | Probable Resistance-Nodulation-Cell Division (RND) efflux transporter | Defence mechanisms –<br>Inorganic ion transport and metabolism |
| **General function prediction only** |  |  |  |
| PA0633 |  | Phage tail protein |  |

To further explore the suggested connection between BatR and PVD and HCN production, we grew WT and Δ*batR* strains in the presence of SIC of the antibiotics PIP, CIP, and GENT, and quantified PVD and HCN production. Contrary to expectations, PVD production was significantly increased in the Δ*batR* strain under PIP challenge in liquid media (Fig. 6a). Likewise, exposure of WT and Δ*batR* strains to Feigl Anger solution, showed enhanced HCN production in the Δ*batR* strain when challenged with PIP (Fig. 6d). These results confirm the link between BatR and both phenotypes but suggest that the association may be highly dependent on growth conditions. Consistent with results from the EVPL assays, we did not observe a significant increase in siderophore production under CIP or GENT challenge conditions in the Δ*batR* strain (Fig. 6 b,c). These findings suggest that, while *batR* deletion leads to a general antibiotic/biofilm tolerance phenotype, some of the specific proteomic changes observed are antibiotic-specific.

**FIG 6.**
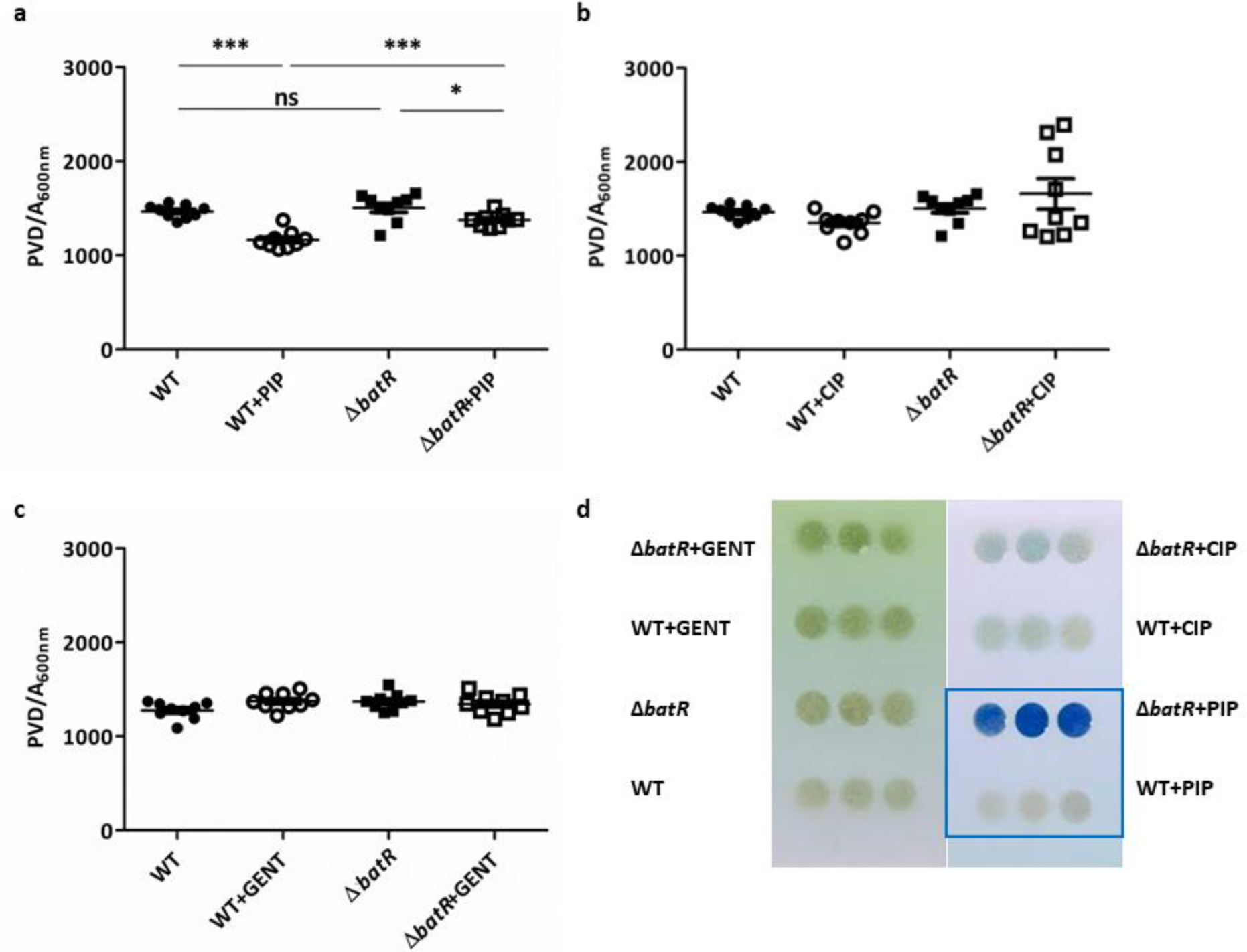
a, b, c. Pyoverdine production. Fluorescence (PVD) relative to growth (A_600_) of WT and Δ*batR* PAO1 cells in liquid medium with and without PIP (a), CIP (b) and GENT (c). PVD: relative fluorescence measured at 460 nm (excitation: 400 nm) for the strains at 65 h. **d. HCN production.** Feigl-Anger assay showing release of HCN from WT and Δ*batR* PAO1 strains. Three independent biological replicates are shown. The blue pigmentation corresponds to HCN release; note the increased intensity of blue colour in the Δ*batR* PAO1 strain when challenged with PIP (blue boxed).

### BatR interacts with two kinase enzymes

To understand the molecular basis of BatR function, we next investigated its interactions with other PAO1 proteins by performing a co-immunoprecipitation (Co-IP) analysis (S2 Data). M2-tagged BatR pulled down several cytoplasmic proteins, indicating potential direct regulatory mechanisms (Table 6). Independent validation of strongly co-precipitating proteins was conducted using Bacterial Two-Hybrid (B2H) analysis. This confirmed specific interactions between BatR and two structurally diverse kinase enzymes: <u>Gl</u>ycerol <u>p</u>hosphate <u>k</u>inase (<u>GlpK</u>) and <u>S</u>tress response <u>k</u>inase A (<u>SrkA</u>) (Fig. 7a). These interactions are noteworthy given the recently established roles of both proteins in antimicrobial tolerance and stress response (33, Liu Y, 2022 #12, 34).

**FIG 7.**
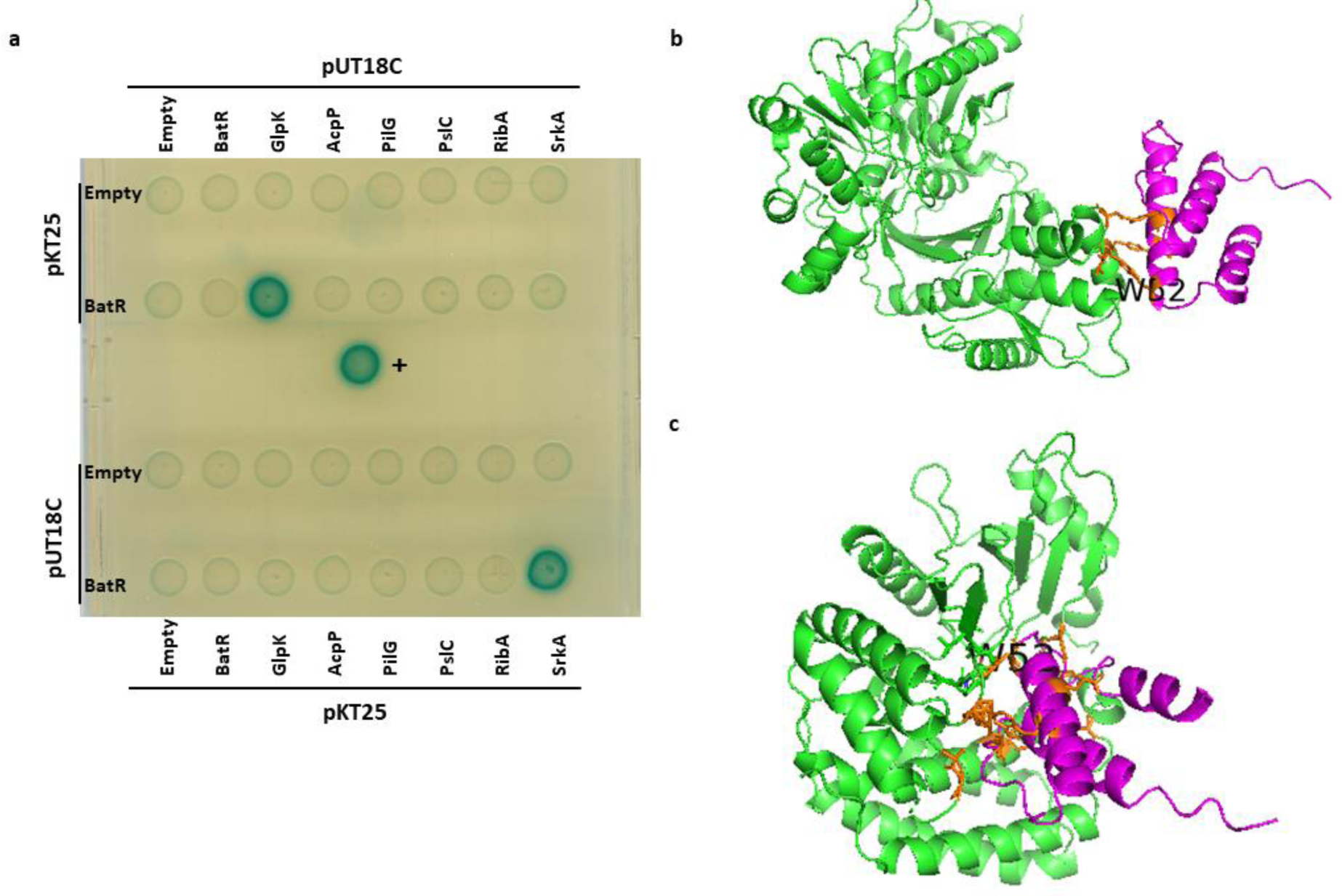
**BatR interacts with two kinase-like proteins. a**. Representative image of qualitative β-galactosidase assays on agar plates. pKT25 and pUT18C fusions are shown in rows and columns, with the indicated protein/empty vector present in each case. Positive control (+): pKT25-zip and pUT18C-zip encoding the two adenylate cyclase fragments, T25 and T18, each fused to the leucine zipper domain of the yeast transcriptional activator GCN4. **b** and **c** are cartoon representations of BatR (magenta) modelled onto the proteins GlpK of SrkA (35), respectively. The interacting residues are shown in yellow and the W_52_ residue is labelled.

**Table 6.**
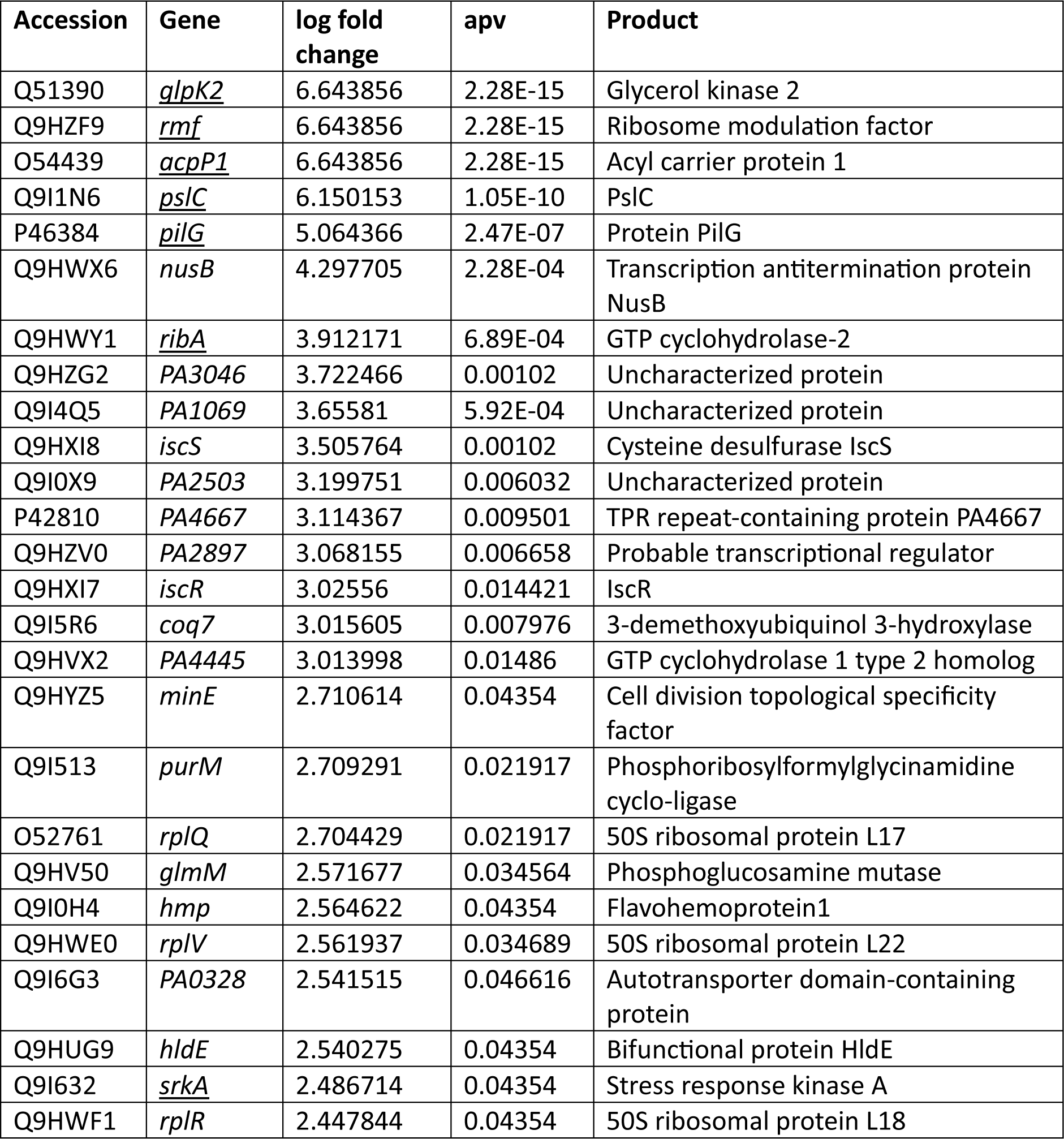
BatR interacting proteins.

To gain additional insight into the potential interaction between BatR and GlpK / SrkA, we used AlphaFold 2 (35) to create three dimensional models of all three proteins. These models allowed us to dock BatR onto GlpK and SrkA, predicting potential interaction residues and providing valuable structural information. Interestingly, the BatR residue W_52_ was common to both predicted interaction interfaces (Fig. 7 b,c). This residue is highly conserved among *Pseudomonas* BatR homologs but is missing from *E. coli* RMF (Fig. S6).

#### GlpK (PA3582)

is an ATP-dependent glycerol kinase that catalyses the first step in the metabolism of glycerol, producing glycerol 3-phosphate (G3P) during aerobic microbial metabolism in *P. aeruginosa* (36). G3P accumulation is associated with reduced cell growth, diminished pyocyanin production, lowered tolerance to oxidative stress and increased kanamycin susceptibility in *P. aeruginosa* (34). The observed phenotypes in the Δ*batR* strain align with those linked to G3P accumulation (Figures 2, 3 & 5), suggesting a potential role for BatR in suppressing GlpK activity. The interaction between BatR and GlpK is predicted to be electrostatic, with the amino acids from BatR (W_44_, E_46_ and W_52_) and GlpK (V_326_, N_328_ and Y_335_) being involved (Fig. 7b). We propose that BatR’s binding suppresses GlpK activity as the binding may interfere with the ATP/ADP predicted binding site (residues 314 & 318, Table 7). Consistent with the expected consequences of G3P accumulation, *batR* deletion significantly affected PAO1 growth and survival when cultured in M9 containing succinate & glycerol, as previously described (34) (Fig. 8a).

**FIG 8.**
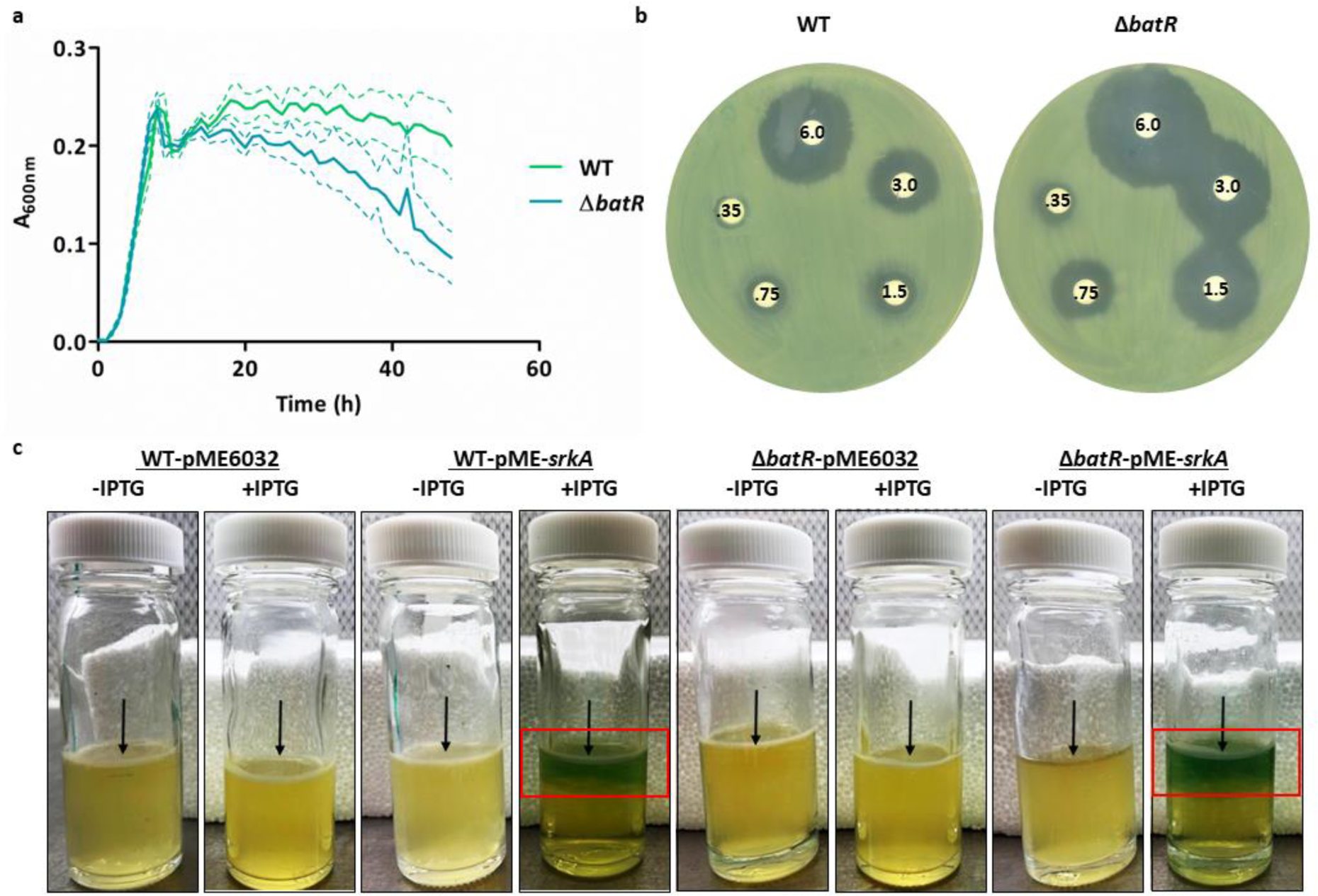
BatR interacts with <u>Gl</u>ycerol <u>p</u>hosphate <u>k</u>inase (GlpK) and <u>S</u>tress <u>r</u>esponse <u>k</u>inase A (<u>SrkA</u>). a. Growth of *P. aeruginosa* PAO1 strains with 20 mM succinate and 20 mM glycerol as the carbon sources. The mean growth for 3 biological replicates for strains WT (35) and Δ*batR* (blue) is shown as a solid line and standard deviation shown as dotted lines. Cells were grown for 48 h at 37°C under static conditions. **b. Hydrogen peroxide (H_2_O_2_) sensitivity assay.** Photographs of the bacterial culture plates 1 d after incubation at 37°C showing halos corresponding to inhibition zones of bacterial growth around the H_2_O_2_ disks, indicating bacterial sensitivity. Phenotypes of the WT and Δ*batR* (highly sensitive) strains are shown. The concentration of H_2_O_2_ on each disk is shown (0.35-6.0 %). **c. Biofilm pellicle assay showing 1 d biofilm and pyocyanin production.** Photograph of static cultures of the WT and the Δ*batR* strains overexpressing *srkA* grown without (-) and with (+) 0.05 mM IPTG demonstrating mature biofilm at air-liquid interface. Two biofilm characteristics were observed: pellicle (arrowed) and pyocyanin production (blue coloration; red boxed).

**Table 7.**
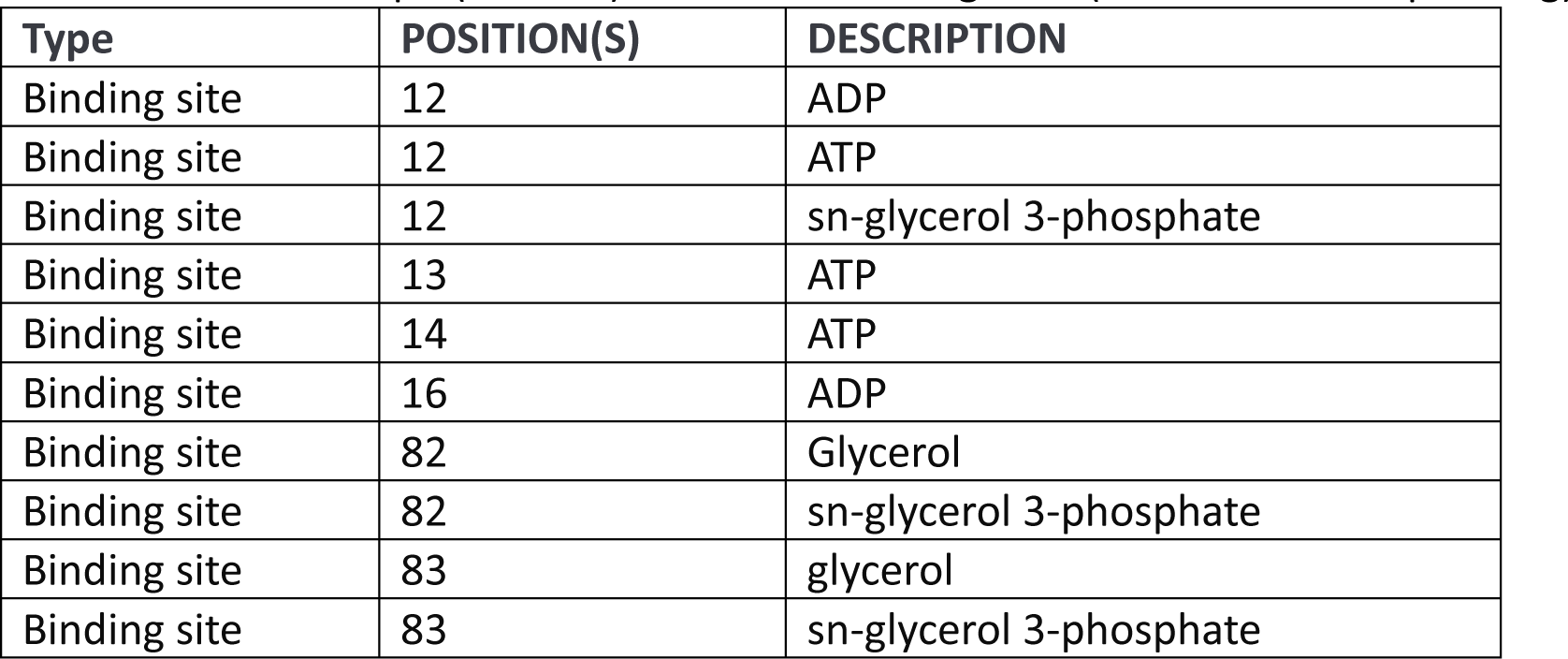
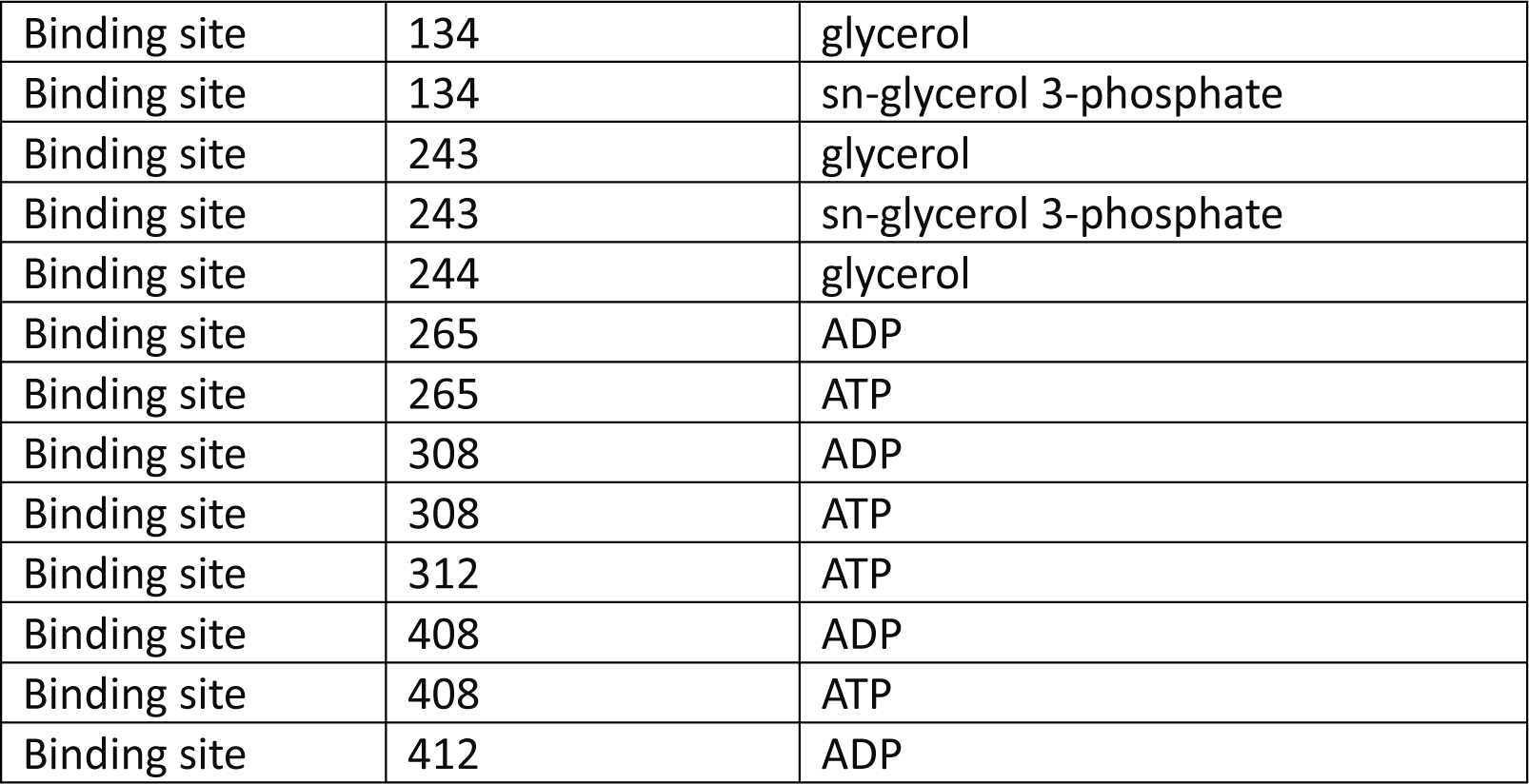
Predicted GlpK (PA3582) active and binding sites (source www.uniprot.org)

#### SrkA (PA0486)

is a eukaryotic-like serine-threonine protein kinase, whose protective role in antimicrobial and environmental stress has been characterised in *E. coli* (33, 37). This protein is linked to a reactive oxygen species (ROS) cascade, and a deficiency of *srkA* stimulates stress-induced <u>p</u>rogrammed <u>c</u>ell <u>d</u>eath (PCD) even after stress dissipated. The deletion of *batR* in *P*. *aeruginosa* strain PAO1 leads to reduced survival rates following antibiotic treatment in biofilms (Figure 2&3), despite no observed changes in MIC values (Table 1), consistent with SrkA dysregulation in *E. coli* (33). Additionally, BatR exhibits a protective role against hydrogen peroxide (H_2_O_2_) stress in PAO1 (Fig. 8b), like SrkA in *E. coli*, supporting a potential connection between BatR and SrkA in stress response pathways. The interaction between BatR and SrkA is also predicted to be electrostatic, involving specific amino acid residues from both BatR (R_26_, W_44_, E_46_ and W_52_) and SrkA (N_32_, Y_34_, P_113_, A_234_, G_235_, Y_276_ and F_293_) (Fig. 7c). We propose that the binding of BatR suppresses the kinase activity of SrkA, possibly impacting its role in ATP binding (residue 33 predicted to bind ATP, see Table 8). This interaction might enhance cellular resistance to stress. To test this hypothesis, we overexpressed *srkA* in the WT and Δ*batR* strains. The overproduction of SrkA in both strains resulted in increased lethality and enhanced pyocyanin production in pellicle biofilms grown in liquid medium (Figure 8c). It is noteworthy that pyocyanin production occurs underneath growing pellicles when *srkA* is overexpressed. Given the lack of characterisation of *srkA* in *P. aeruginosa* so far, we propose a potential role in regulating antimicrobial stress and pyocyanin production, two phenotypes associated with *batR* in this study.

**Table 8.**
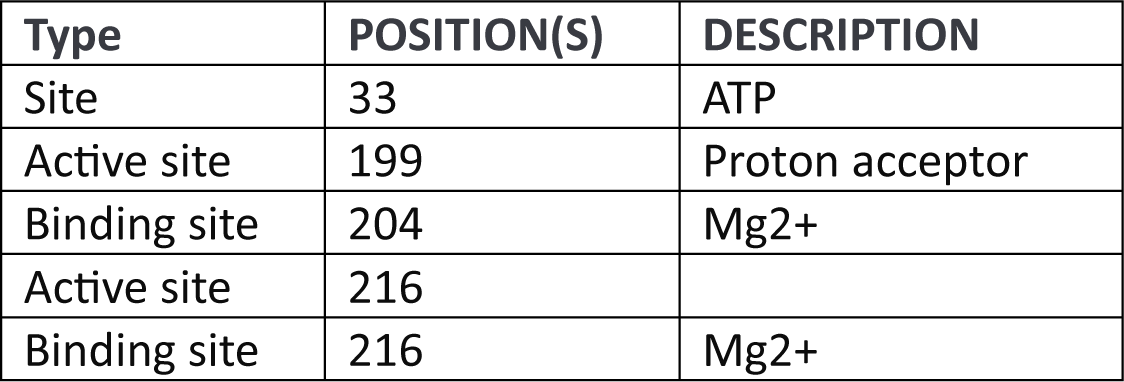
Predicted SrkA (PA0486) active and binding sites (source www.uniprot.org)

## DISCUSSION

In this study, we characterise the *P. aeruginosa* hydrophilin BatR and determine its role in antimicrobial tolerance and biofilm formation, and its potential clinical significance. *batR* homologs exhibit a high degree of conservation within diverse bacteria, particularly in ɣ-proteobacterial species (23). The identification of a well-supported clade of Pseudomonadales carrying *batR* homologs that is both genetically and structurally distinct from the characterised Enterobacterales/Vibrionales *rmf* clade suggests a divergent evolutionary path for these genes, and a distinct cellular function for BatR. The well characterised *batR* homolog in *E. coli*; *rmf*, contributes to ribosome hibernation and tolerance to the aminoglycoside antibiotics gentamicin and netilmicin (20, 38). Consistent with this, the binding site of this hibernation factor is near to those described for these antibiotics, thus potentially interfering with its mechanism of action (23). Conversely, the contribution of *batR* to antimicrobial tolerance is not linked to antimicrobial mode-of-action, making it highly unlikely that BatR functions solely via ribosomal inhibition, as previously suggested.

BatR contributes to both biofilm formation and antibiotic tolerance in conditions like those in CF infections. Our data suggest that BatR is particularly important during the early stages of *P. aeruginosa* lung tissue infection at sub-inhibitory concentrations of antibiotics. This is relevant in CF patients since they are typically subject to extended antibiotic regimes, but the drugs do not necessarily reach the entire lung at inhibitory concentrations (39). Both WT and Δ*batR* strains formed sponge-like biofilm structures, characteristic of CF infections after 7 days PI in the EVPL model. However, upon challenge with CIP the Δ*batR* strain formed a less resistant biofilm structure on the surface of the EVPL bronchiolar tissue compared to the WT strain. Interestingly, whilst these differences were visually obvious, similar CFU were recovered from tissue infected by both genotypes at 7 days. This is consistent with a large fraction of dead or unculturable cells in WT biofilms, as previously observed for *P. aeruginosa* PA14 biofilms (27). Noteworthy, transcriptomic analysis of *P. aeruginosa* strain PA14 in EVPL model revealed significant differential expression of *batR* at 7 days compared with 1 day PI (40). BatR is also involved in *P. aeruginosa* biofilm tolerance in the synthetic chronic wound model of diabetic foot infections. This is consistent with the finding that *batR* expression is differentially increased in burn wound infections (41), and underscores the versatility of *batR* in mediating *P. aeruginosa* pathogenesis across various infection settings.

BatR function appears to be consistently associated with pyocyanin production. This molecule was notably reduced in Δ*batR* following exposure to CIP in the EVPL model (Figure 5). Acting as a potent electron acceptor, pyocyanin influences cellular redox balance, inducing oxidative stress in host cells and ultimately leading to cell damage and lysis (42). Within the oxygen-limited environment of *P. aeruginosa* biofilms, pyocyanin is crucial for metabolic continuity and significantly impacts the biofilm’s response to antibiotic treatments (43–45). Our proteomic results align with previous research demonstrating the induction of denitrification enzymes by phenazine deficiency in *P. aeruginosa* biofilms (46). Additionally, upregulation of the permease FeoB (Table X) by *batR* facilitates efficient iron uptake in biofilms, highlighting the intricate interplay between BatR, phenazine metabolism and iron homeostasis (47). Consistent with our H&E staining results (Figure 4), pyocyanin production has been linked to biofilm architecture and eDNA production in *P. aeruginosa* (48), contributing to the observed differences in biofilm structure.

Finally, to understand the molecular basis of BatR function we investigated its protein interaction partners within the cell. Through screening, two proteins; SrkA and GlpK, were identified. BatR interaction with SrkA and GlpK control antimicrobial tolerance and virulence factor production of *P. aeruginosa* PAO1 biofilms. Thus, BatR may have potential therapeutic utility as a target for the control of *P. aeruginosa* infections. Similarly to *E. coli*, SrkA in *P. aeruginosa* may have a regulatory role in stress mediated PCD, mediated by BatR interaction. *E.coli* SrkA is partially regulated by the Cpx envelope stress-response system, which has both protective and destructive roles that help bacteria make a live-or-die decision in response to stress (33). Interestingly, transcription of *rmf* is also induced by the Cpx system in *E. coli*, suggesting a link between them (49). Our results also suggest the involvement of SrkA in regulating pyocyanin production in *P. aeruginosa*, although further research is necessary to fully understand this link.

G3P metabolism has been characterized in *P. aeruginosa* due to its relevance to CF infections (34): glycerol is released from the cleavage of phosphatidylcholine, the major lung surfactant in CF patients (50). G3P is involved in maintaining cellular homeostasis, and increased levels of G3P lead to reduced pyocyanin production and resistance to kanamycin (34). This represents a clear example of bacterial antibiotic resistance closely correlated with physiological metabolism (51, 52). However, the specific mechanism by which G3P accumulation induces phenotypic alterations in *P. aeruginosa* PAO1 remains unclear.

Our current view of the mechanism of action of this novel post-transcriptional regulator, BatR, expressed under biofilm conditions that plays a role under antimicrobial stress in the opportunistic human pathogen *P. aeruginosa,* is sketched in Fig. 9.

**FIG 9.**
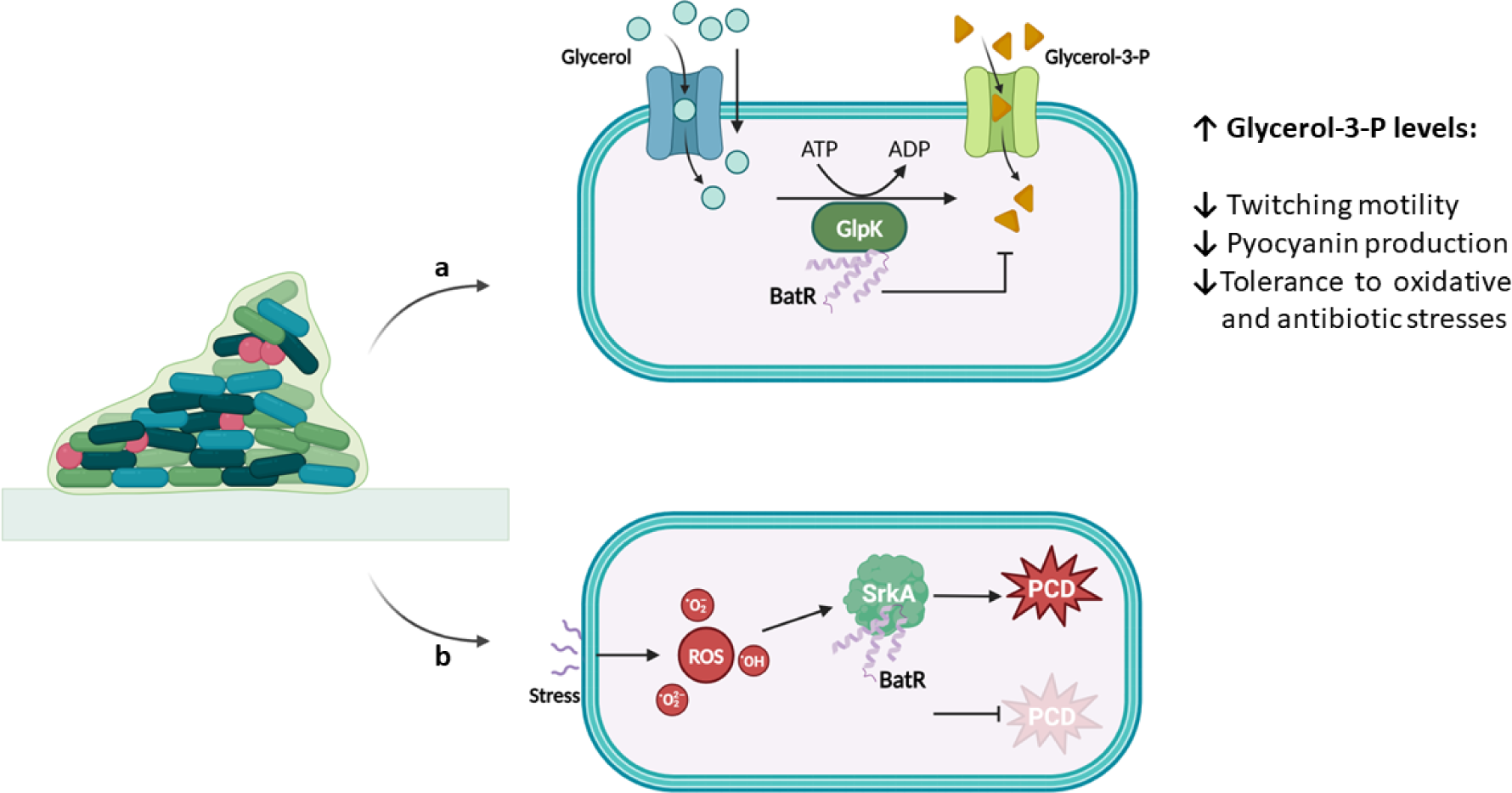
Schematic representation of BatR mechanism of action. a. BatR-GlpK interaction and its impact on G3P metabolism in *P. aeruginosa* PAO1. G3P, crucial for various cellular processes, can be imported from the extracellular environment or derived from glycerol phosphorylation via GlpK activity. The accumulation of G3P reduces antibiotic resistance, pyocyanin production, oxidative stress tolerance, and twitching motility. GlpK activity is suppressed upon interaction with BatR, potentially resulting in decreased G3P levels and increased pyocyanin and antimicrobial resistance. **b**. **BatR-SrkA interaction and its implications in stress-mediated bacterial PCD and pyocyanin production**: SrkA, plays a protective role against stressors such as H_2_O_2_ and antibiotics, controls pyocyanin production, and interacts with BatR. This interaction is hypothesised to be crucial for SrkA’s protective function and suggests a novel mechanism for stress response modulation in *P. aeruginosa*.

In conclusion, our findings expand the understanding of molecular mechanisms contributing to antimicrobial tolerance in *P. aeruginosa* biofilms. These results have broad implications for the functions of uncharacterised proteins induced under biofilm conditions, shedding light on their pivotal role in orchestrating multifaceted processes. This deeper insight enhances our ability to develop targeted therapeutic interventions to combat biofilm-associated challenges.

## MATERIALS AND METHODS

### Bioinformatic analysis

A phylogenetic tree of RMF proteins was constructed using 765 publicly available protein sequences from the NCBI database. The dataset was curated based on a criterion of 50% sequence identity and 40% query cover to ensure the representation of diverse homologs while maintaining a reasonable level of similarity. The multiple sequence alignment was performed using Clustal Omega (v1.2.4), generating an alignment matrix with 137 columns and 135 distinct patterns. Phylogenetic signal analysis revealed 82 parsimony-informative sites, 33 singleton sites, and 22 constant sites. Manual curation was done to remove the repetitive sequences and false hits. The phylogeny estimation was done using IQ-TREE, (multicore v1.6.12), employing the maximum likelihood (ML) criterion. The tree visualization was done using iTOL (v6.8.1), representation chosen was an unrooted tree with branch lengths proportional to the inferred evolutionary distances between sequences. The clade colours were assigned at order level of the taxonomy.

### Bacterial strains and growth media

Bacterial strains and plasmids used in this study are listed in Table 9. Unless otherwise stated, *P. aeruginosa* PAO1 and *E. coli* DH5α strains were routinely cultured in lysogeny broth (LB (JH, 1972 #6)) at 37°°C solidified with 1.5% w/v agar where appropriate. For Gly + Succinate growth curves, PAO1 strains were grown in M9 salts supplemented with 1 mM MgSO_4_, 1 mM CaCl_2_ (53), adding 20 mM succinate and 20 mM glycerol as the carbon sources. For growth curves, measurements were taken every 30 min for up to 48 h on a FLUOstar nano plate reader (BMG) with the plate being incubated at 37°°C under static or planktonic conditions, as indicated.

**Table 9.** Strains and plasmids used in this study.

| Strain | Description | Reference |
| --- | --- | --- |
| ***P. aeruginosa*** |  |  |
| WT PAO1 | Wild-type *Pseudomonas aeruginosa* PAO1 | 1 |
| $\Delta batR$ PAO1 | Non-polar deletion of *PA3049 (batR)* | This study |
| ***E. coli*** |  |  |
| DH5α | *endA1*, *hsdR17*(r<sub>K</sub>-m<sub>K</sub><sup>+</sup>), *supE44*, *recA1*, *gyrA* (Nal<sup>r</sup>), *relA1*, $\Delta(lacIZYA-argF)$ U169, *deoR*, $\Phi 80dlac\Delta(lacZ)M15$ | 2 |
| BTH101 | *F*<sup>-</sup>, *cya-99*, *araD139*, *galE15*, *galK16*, *rpsL1 (Strr)*, *hsdR2*, *mcrA1*, *mcrB1*. | 3 |
| **Plasmids** |  |  |
| pTS1 | Tet<sup>R</sup>, suicide vector; *ColE1*-replicon, *IncP-1*, *Mob*, *lacZ* | 4 |
| pTS1-*batR* vector | pTS1 with $\Delta batR$ constructs as BamHI-HindIII inserts | This study |
| pME6032 | Tet<sup>R</sup>, P<sub>K</sub>, 9.8 kb pVS1 derived shuttle vector | 5 |
| pME-*batR* | pME6032 containing the ORF of *batR* between the EcoRI-XhoI restriction sites | This study |
| pME-3xFLAG-*batR* | pME6032 containing the 3x-Flag-tagged BatR ORF using EcoRI-XhoI restriction sites encoded in the outer primers (used in Co-IP assay) |  |
| pKT25 | Plasmid for constructing N-terminal fusions to T25, Kan<sup>R</sup> | 3 |
| pKT25-BatR | pKT25 with the ORF of *batR* cloned within the XbaI/KpnI sites (used in B2H assay) | This study |
| pKT25-AcpP | pKT25 with the ORF of *acpP* (PA2966) cloned within the XbaI/KpnI sites (used in B2H assay) | This study |
| pKT25-PslC | pKT25 with the ORF of *pslC* (PA2233) cloned within the XbaI/KpnI sites (used in B2H assay) | This study |
| pKT25-PilG | pKT25 with the ORF of *pilG* (PA0408) cloned within the XbaI/KpnI sites (used in B2H assay) | This study |
| pKT25-RibA | pKT25 with the ORF of *ribA* (PA4047) cloned within the XbaI/KpnI sites (used in B2H assay) | This study |
| pKT25-SrkA | pKT25 with the ORF of *srkA* (PA0486) cloned within the XbaI/KpnI sites (used in B2H assay) | This study |
| pKT25-GlpK | pKT25 with the ORF of *glpK* (PA3582) cloned within the XbaI/KpnI sites using Gibson assembly (B2H assay) | This study |
| pUT18C | Plasmid for constructing C-terminal fusions to T18, Carb<sup>R</sup> | 3 |
| pUT18C-BatR | pUT18C with the ORF of *batR* cloned within the XbaI/KpnI sites (used in B2H assay) | This study |
| pUT18C-AcpP | pUT18C with the ORF of *acpP1* (PA2966) cloned within the XbaI/KpnI sites (used in B2H assay) | This study |
| pUT18C-PslC | pUT18C with the ORF of *pslC* (PA2233) cloned within the XbaI/KpnI sites (used in B2H assay) | This study |
| pUT18C-PilG | pUT18C with the ORF of *pilG* (PA0408) cloned within the XbaI/KpnI sites (used in B2H assay) | This study |
| pUT18C-RibA | pUT18C with the ORF of *ribA* (PA4047) cloned within the XbaI/KpnI sites (used in B2H assay) | This study |
| pUT18C-SrkA | pUT18C with the ORF of *srkA* (PA0486) cloned within the XbaI/KpnI sites (used in B2H assay) | This study |
| pUT18C-GlpK | pUT18C with the ORF of *glpK* (PA3582) cloned within the XbaI/KpnI sites using Gibson assembly (B2H assay) | This study |
| pKT25-Zip | pKT25 carrying the leucine zipper of GCN4, Kan<sup>R</sup> (used as positive control in B2H assay) | 3 |
| pUT18-zip | pUT18 carrying the leucine zipper of GCN4, Kan<sup>R</sup> (used as positive control in B2H assay) | 3 |

Carbenicillin (Carb) was used at 100 μg/ml, Kanamycin (Kan) at 50 μg/ml, Tetracycline (Tet) at 12.5 μg/ml for *E. coli* and 100 μg/ml for *P. aeruginosa*, IPTG at 0.5 mM, and X-gal at 40 μg/ml. The antibiotics Piperacillin (PIP), Gentamicin (GENT), and Ciprofloxacin (CIP) were employed at concentrations optimized for each experiment, with details provided accordingly.

### Molecular biology techniques and Genetic manipulation of PAO1

These procedures were performed as previously described (54). All pTS1 plasmid inserts were synthesised and cloned into pTS1 by Twist Bioscience.

The ORF of *batR* and *srkA* were amplified by PCR with primers batR_EcoRI_F / batR_XhoI_R and srkA_EcoRI_F / srkA_XhoI_R (Table 10), respectively, and ligated between the EcoRI and XhoI sites of pME6032. For the flag-tagged *batR* construction in pME6032, primers batR_EcoRI_F and 3xflag_batR_XhoI (Table 10) were used. Bacterial-2-hybrid plasmids were made by Gibson assembly (*glpK*) and restriction cloning (*batR, acpP, srkA, pslC, pilG*, and *ribA*) into the XbaI and KpnI sites of pKNT25 and pUT18C using the primers indicated in Table 10.

**Table 10.**
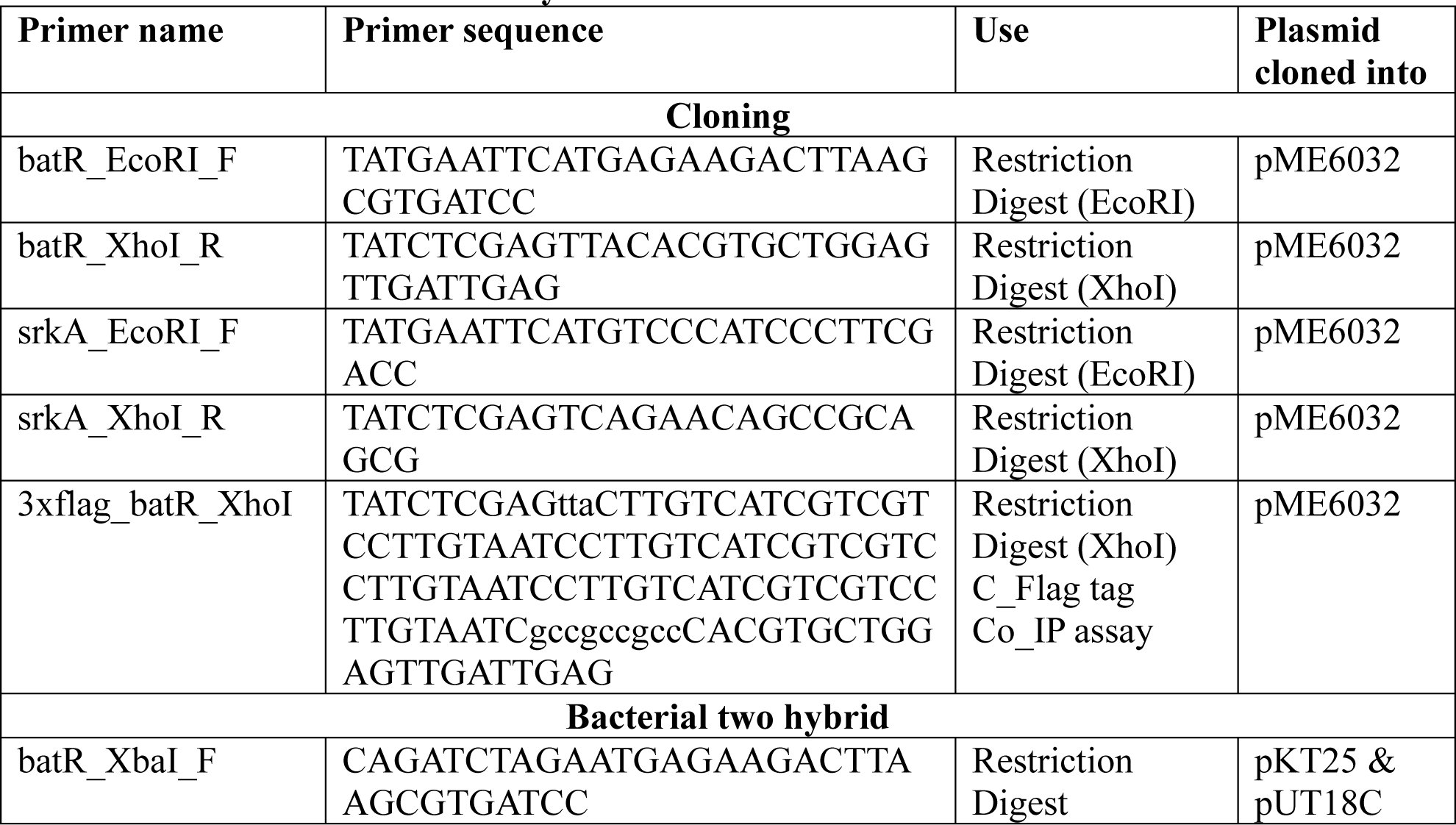
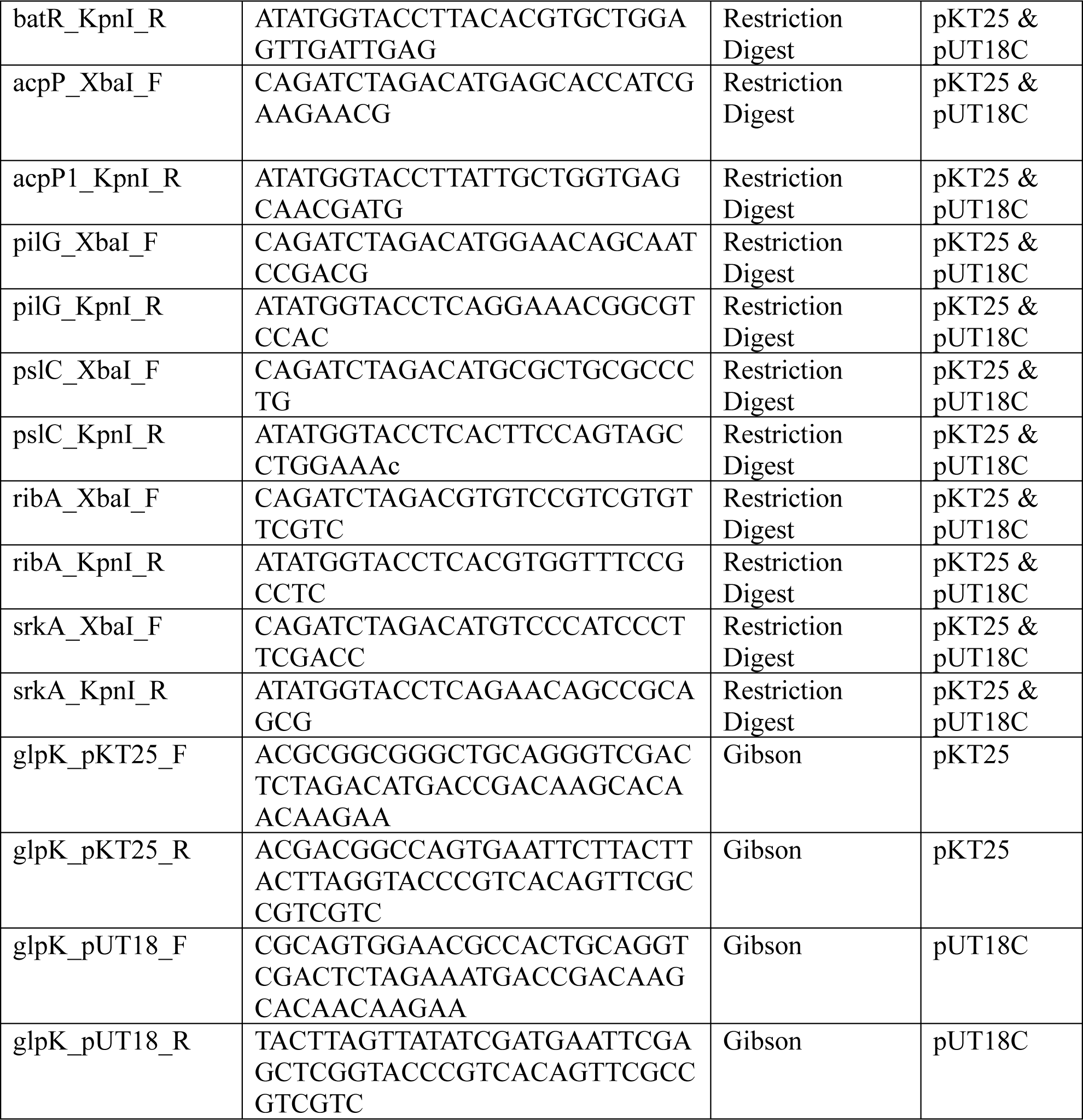
Primers used in this study.

### MIC determination

Minimum inhibition concentrations of antibiotics were determined by the broth microdilution method (24) following the EUCAST guidelines, using Mueller-Hinton broth. The Sub Inhibitory Concentration (SIC) was defined as being 1/4 of the lowest antibiotic concentration that inhibited visible growth after overnight incubation at 37°C.

### Inhibition disc assay

Bacterial cultures were grown in LB medium at 37°C to mid-log phase, A_600nm_=0.5, and 150 µl were spread on each plate. Discs containing the antibiotics/H_2_O_2_ were gently placed on the agar and plates were incubated inverted overnight. The normalized width of the antimicrobial “halo” (NW_halo_) of each disk was determined after (55).

### Glass beads biofilms and Crystal Violet (CV) assays

These assays were performed as described elsewhere (25) with the following modifications. Ten independent biological replicates were included: five PIP-exposed biofilm lineages (challenged with 0.25× MIC of PIP for 90 min) and five unexposed control lineages. Cells recovered from the beads were serial diluted and spotted onto LB plates for CFU counting.

For the CV assay, the A_590nm_ was measured at using a SPECTROstar nano plate reader (BMG Labtech).

### Membrane permeability assays

These assays were performed as described elsewhere (56).

### EVPL infection model

EVPL was prepared as previously described (27, 57). Porcine lungs were obtained from two local butchers (Quigley and Sons, Cubbington; and Taylor’s Butcher, Earlsdon) and dissected on the day of delivery under sterile conditions. Following infection of bronchiolar tissue pieces, 500 μl of SCFM ± 16 μg/ml CIP was added to each well. Tissue pieces were incubated at 37°C for 2 and 7 d. Uninfected controls were included. EVPL biofilm recovery and assessment of bacterial load and virulence factors production were determined as described elsewhere (27).

### Haematoxylin & eosin staining

H & E staining was assayed as previously described (27). The infected/uninfected EVPL tissue pieces were fixed and sent to the University of Manchester Histology Core Facility for paraffin wax embedding, sectioning, and mounting. Samples were de-paraffinized and stained in Mayer’s hemalum solution (Merck Millipore) and counterstained in eosin Y solution (Merck Millipore). Images were taken using a Zeiss Axio Imager Z2 light microscope with the Zeiss AxioCam 506 and Zeiss Zen Blue v2.3 pro software.

### Synthetic chronic wound infection model

These assays were performed as previously described (28).

### Quantitative Proteomics (TMT) for expression analysis

The detailed protocol is presented in the supplemental materials section (S3 Data). The mass spectrometry proteomics data have been deposited to the ProteomeXchange Consortium via the PRIDE (58) partner repository with the dataset identifier PXD050997 and 10.6019/PXD050997.

### Hydrogen cyanide (HCN) production

The Feigl-Anger assay was employed to detect HCN production. These assays were performed as described elsewhere (59).

### Co-Immunoprecipitation and mass spectrometry analysis

The detailed protocol is presented in the supplemental materials section (S4 Data). Data are available via ProteomeXchange with identifier PXD050995.

### Bacterial 2 hybrid assays

These assays were performed as described elsewhere (60) with some modifications. The ORFs of *batR, acpP, pilG, pslC, ribA, srkA* and *glpK* were cloned into pKT25 and pUT18C using either conventional restriction enzyme cloning or Gibson assembly, as indicated in Table 10.

### Data presentation and statistical analyses

All graphs and one-way ANOVA followed by post-hoc Tukey’s Multiple Comparison Test, where appropriate, were performed using GraphPad Prism version 5.04 for Windows, www.graphpad.com.

## ACKNOWLEDGMENTS

The authors would like to Govind Chandra for his feedback on bioinformatic analysis. This work was supported by BBSRC Grants BB/X010996/1, BBS/E/J/000PR9797, and BB/T004363/1. AP’s placement in FH’s lab was supported by the Flexible Talent Mobility Award Scheme (FTMA).

## SUPPLEMENTAL MATERIAL

**FIG S1.**
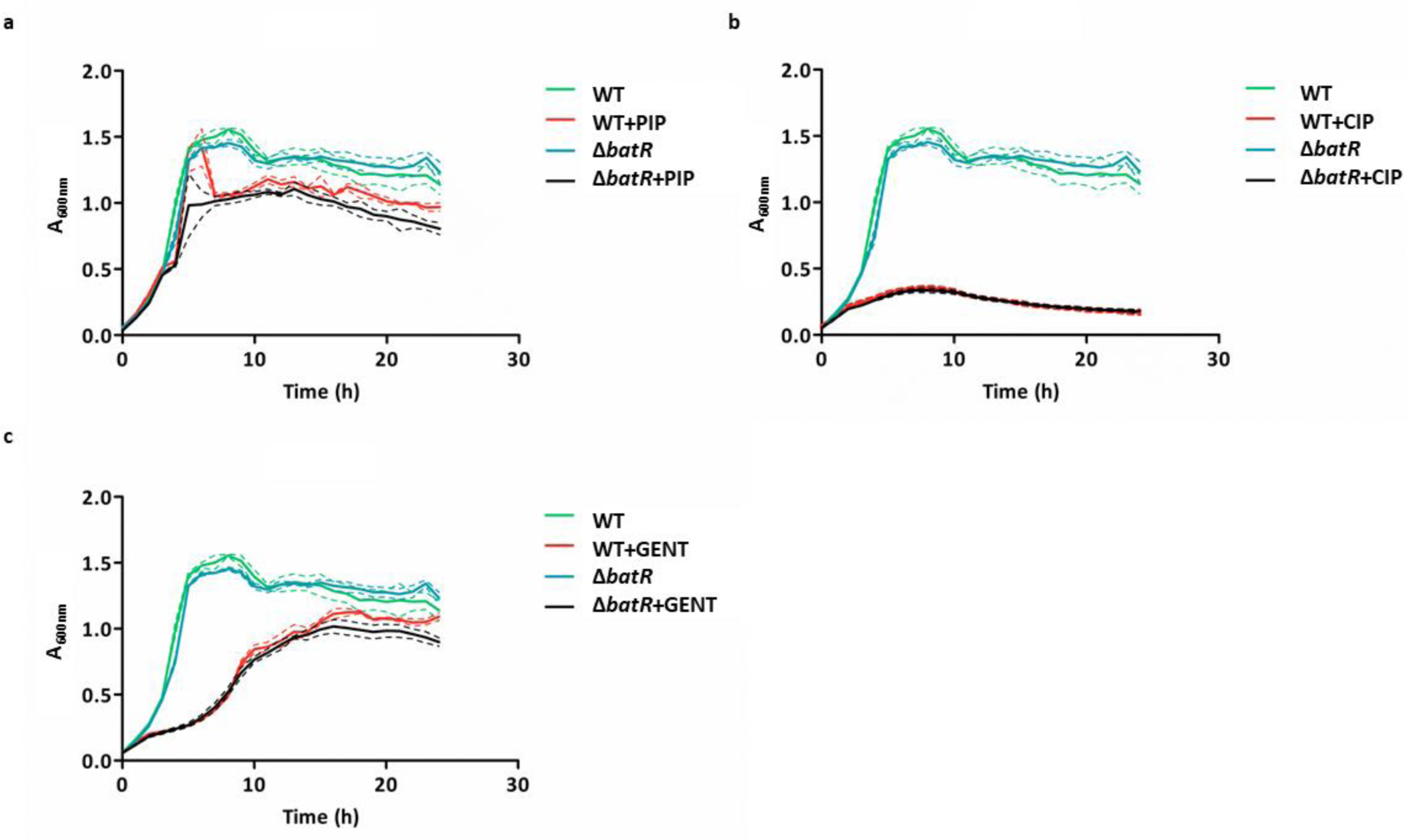
Growth curves are shown for strains WT (35), Δ*batR* (blue) in LB medium and WT (35), Δ*batR* (black) in LB medium supplemented with SIC of CIP, GENT or PIP. The mean growth for 3 biological replicates is shown as a solid line and standard deviation shown as dotted lines. Cells were grown for 24 h at 37⁰C under shaking conditions.

**FIG S2.**
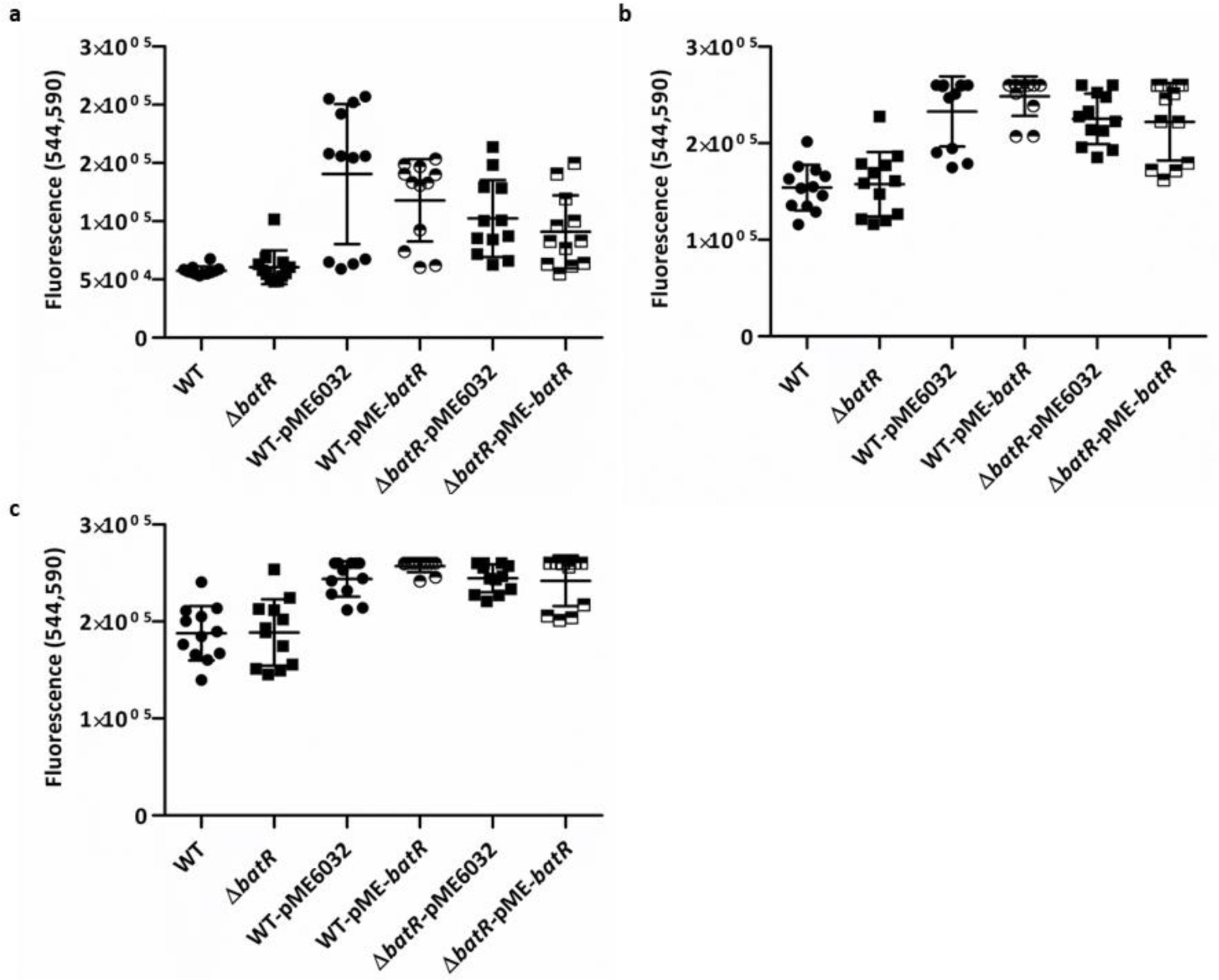
Membrane permeability assays. Drug accumulation as measured by resazurin fluorescence at excitation 544 nm and emission of 590 nm (544,590) is shown. Lines indicate average from 3 biological replicates, with 4 technical replicates for each.

**FIG S3.**
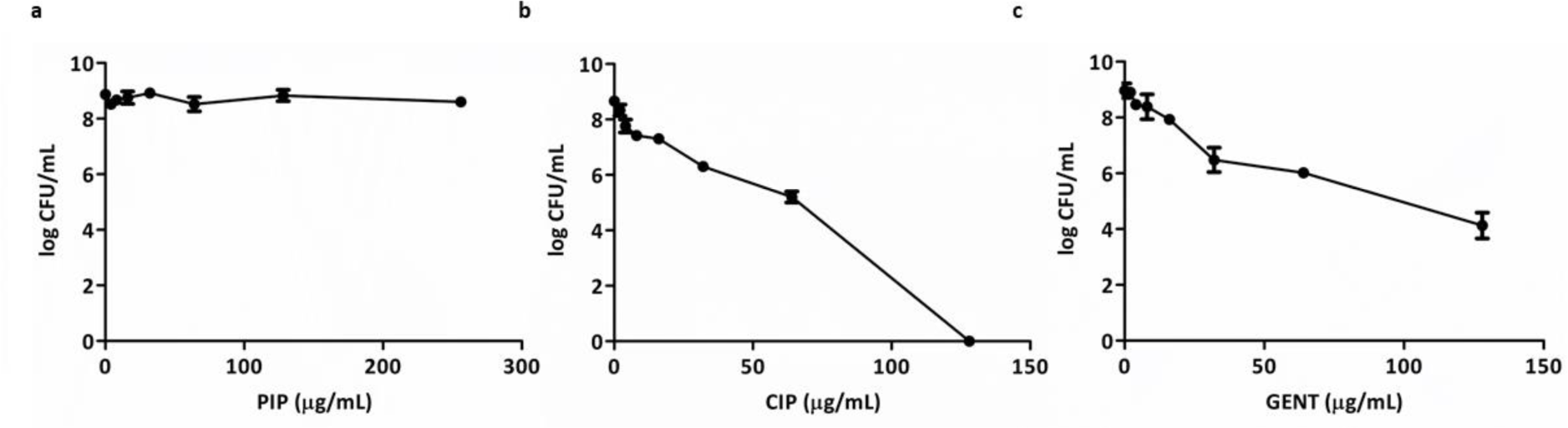
Log of total CFU of *P. aeruginosa* strain PAO1 WT recovered from the EVPL model following treatment with antibiotics. Each strain was grown on EVPL tissue for 48 h then transferred to antibiotic or PBS as a control for 18 h and the CFU/lung determined.

**FIG S4.**
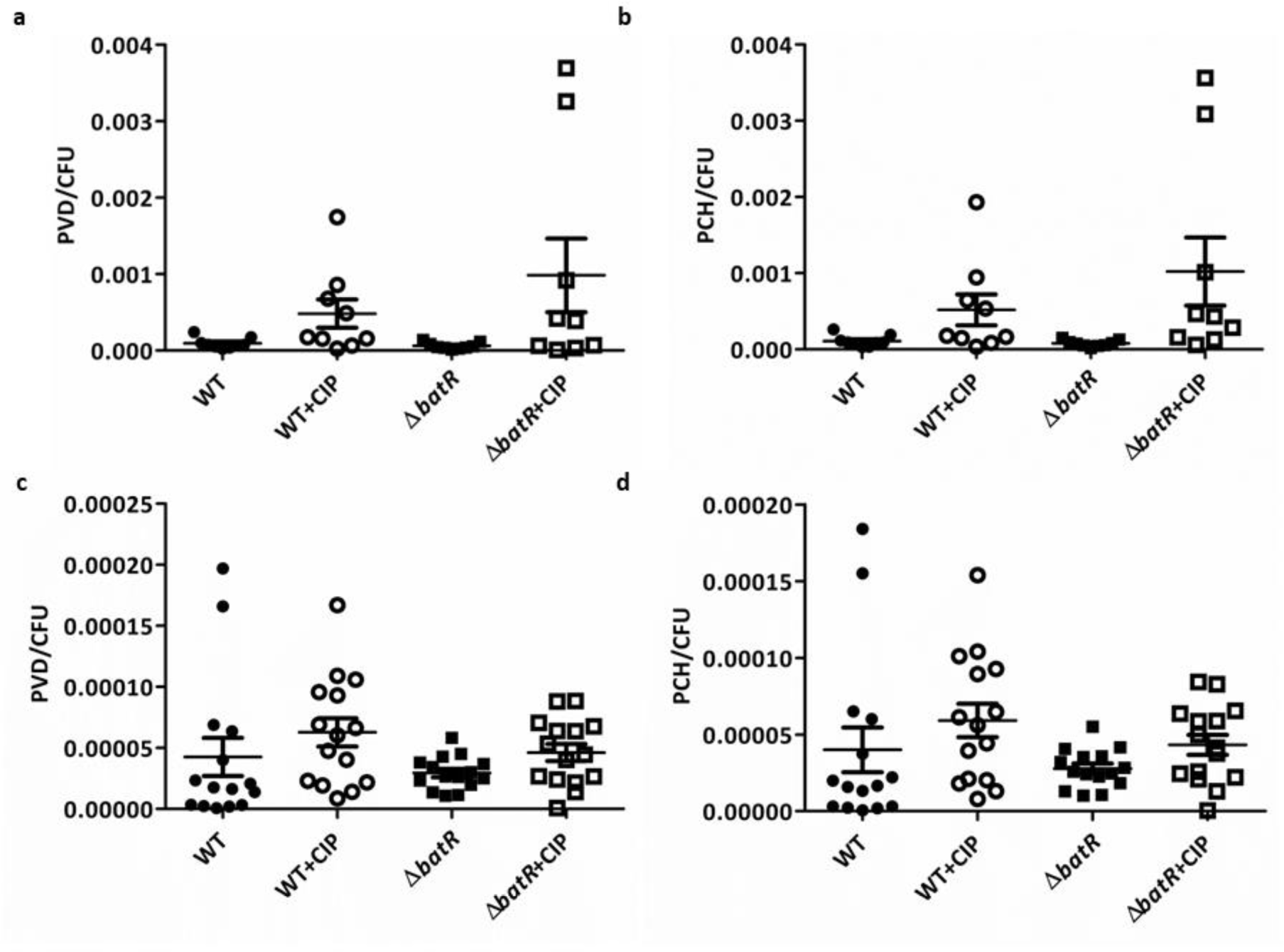
Production of siderophores by *P. aeruginosa* WT and Δ*batR* biofilms in the EVPL model. Units are as follows: for pyoverdine (PVD), fluorescence with excitation/emission of 400±20/460±20 nm per CFU and pyochelin (PCH), fluorescence with excitation/emission of 360±35/460±20 nm per CFU of *P. aeruginosa*. Graphs **a** and **c** show PVD production at 2 d and 7 d PI, respectively. Graphs **b** and **d** show PCH production at 2 d and 7 d PI, respectively. The bars denote means and an ANOVA test shows non-significant differences.

**FIG S5.**
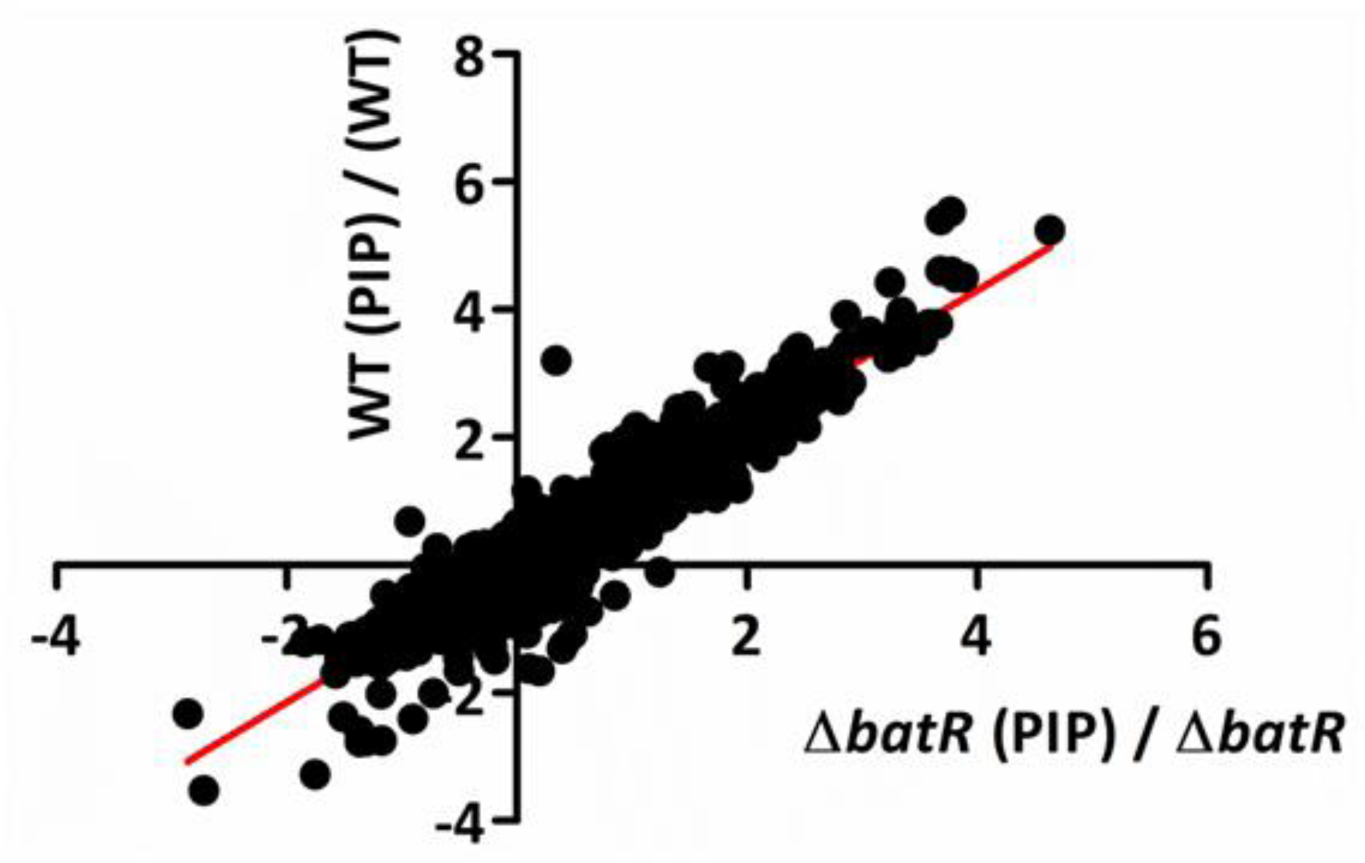
Proteomic analysis: Scatterplot representing pairwise comparison of mean log2 protein abundance values for *P. aeruginosa* WT and Δ*batR*, with and without PIP challenge. A complete list of genes and information on their predicted functions are given in Table 2 and Table 3.

**FIG S6.**
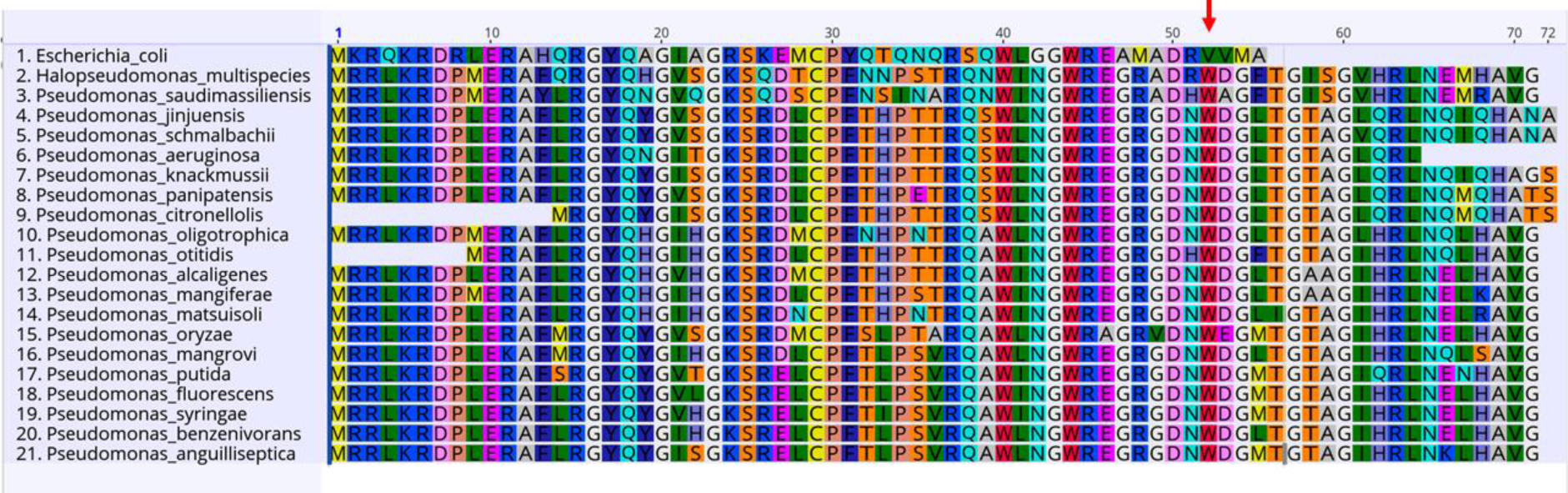
Multiple sequence alignment of selected BatR homologs. ClustalW was used to align RMF of *E. coli* and BatR homologs of different *Pseudomonas* species. The sequence alignment was visualized using Geneious software. The conserved tryptophan residue at position 52 in *Pseudomonas* spp. is shown (arrowed).

**S1 Data.** Integrated proteomic data comparing PAO1 WT and Δ*batR* (±PIP). Contains underlying data for Fig S5. (XLSX).

**S2 Data.** Co-IP data comparing PAO1 WT-pME-*batR* and pME-3xFLAG-*batR*. Contains underlying data for Table 6. (XLSX).

**S3 Data.** Protocol used for Quantitative Proteomics (TMT) for expression analysis. (DOCX)

**S4 Data.** Protocol used for Co-Immunoprecipitation and mass spectrometry analysis. (DOCX)

## REFERENCES

1. Rice LB. 2008. Federal funding for the study of antimicrobial resistance in nosocomial pathogens: no ESKAPE. J Infect Dis 197:1079–81.

2. Shigemura K, Arakawa S, Sakai Y, Kinoshita S, Tanaka K, Fujisawa M. 2006. Complicated urinary tract infection caused by *Pseudomonas aeruginosa* in a single institution (1999–2003). International Journal of Urology 13:538–542.

3. Chitkara YK, Feierabend TC. 1981. Endogenous and exogenous infection with *Pseudomonas aeruginosa* in a burns unit. Int Surg 66:237–40.

4. Mulcahy LR IV, Lewis K. 2014. Pseudomonas aeruginosa biofilms in disease. Microb Ecol doi:10.1007/s00248-013-0297-x.

5. Kielhofner M, Atmar RL, Hamill RJ, Musher DM. 1992. Life-Threatening *Pseudomonas aeruginosa* Infections in Patients with Human Immunodeficiency Virus Infection. Clinical Infectious Diseases 14:403–411.

6. Cassini A, Hogberg LD, Plachouras D, Quattrocchi A, Hoxha A, Simonsen GS, Colomb-Cotinat M, Kretzschmar ME, Devleesschauwer B, Cecchini M, Ouakrim DA, Oliveira TC, Struelens MJ, Suetens C, Monnet DL, Burden of AMRCG. 2019. Attributable deaths and disability-adjusted life-years caused by infections with antibiotic-resistant bacteria in the EU and the European Economic Area in 2015: a population-level modelling analysis. Lancet Infect Dis 19:56–66.

7. ECDPC. 2019. Surveillance of antimicrobial resistance in Europe 2018 doi:10.2900/22212. European Centre for Disease Prevention and Control, Stockholm.

8. Lyczak JB CC, Pier GB. 2002. Lung infections associated with cystic fibrosis. Clin Microbiol Rev doi: 10.1128/CMR.15.2.194-222.2002.

9. Pressler T, Bohmova C, Conway S, Dumcius S, Hjelte L, Høiby N, Kollberg H, Tümmler B, Vavrova V. 2011. Chronic *Pseudomonas aeruginosa* infection definition: EuroCareCF Working Group report. Journal of Cystic Fibrosis 10:S75–S78.

10. Boyd CD OTG. 2012. Second messenger regulation of biofilm formation: breakthroughs in understanding c-di-GMP effector systems. Annu Rev Cell Dev Biol doi:10.1146/annurev-cellbio-101011-155705.

11. Serra DO HR. 2014 Stress responses go three dimensional - the spatial order of physiological differentiation in bacterial macrocolony biofilms. Environ Microbiol doi:10.1111/1462-2920.12483.

12. Piazza A PL, Ciancio Casalini L, Sisti F, Fernández J, Malone JG, Ottado J, Serra DO, Gottig N. 2022. Cyclic di-GMP Signaling Links Biofilm Formation and Mn(II) Oxidation in Pseudomonas resinovorans. mBio 10.1128/mbio.02734-22.

13. Williamson KS RL, Perez-Osorio AC, Pitts B, McInnerney K, Stewart PS FM. 2012. Heterogeneity in Pseudomonas aeruginosa biofilms includes expression of ribosome hibernation factors in the antibiotic tolerant subpopulation and hypoxia-induced stress response in the metabolically active population. J Bacteriol 194:2062–73.

14. Crabbé A JP, Bjarnsholt T, Coenye T. 2019. Antimicrobial Tolerance and Metabolic Adaptations in Microbial Biofilms. Trends Microbiol doi:10.1016/j.tim.2019.05.003.

15. Schobert M JD. 2010. Anaerobic physiology of Pseudomonas aeruginosa in the cystic fibrosis lung. International Journal of Medical Microbiology 300:549–556.

16. Schurek KN MA, Taylor PK, Wiegand I, Semenec L, Khaira BK, Hancock RE. 2008. Novel genetic determinants of low-level aminoglycoside resistance in Pseudomonas aeruginosa. Antimicrob Agents Chemother doi:10.1128/AAC.00507-08.

17. Kaleta MF PO, Zampaloni C, Garcia-Alcalde F, Parker M & Sauer K. 2022. A previously uncharacterized gene, PA2146, contributes to biofilm formation and drug tolerance across the ɣ-Proteobacteria. npj Biofilms and Microbiomes 8:54.

18. Garay-Arroyo A C-FJ, Garciarrubio A, Covarrubias AA. 2000. Highly hydrophilic proteins in prokaryotes and eukaryotes are common during conditions of water deficit. J Biol Chem doi:10.1074/jbc.275.8.5668.

19. Yamagishi M MH, Wada A, Sakagami M, Fujita N, Ishihama A. 1993. Regulation of the Escherichia coli rmf gene encoding the ribosome modulation factor: growth phase- and growth rate-dependent control. EMBO J doi:10.1002/j.1460-2075.1993.tb05695.x.

20. McKay SL PD. 2015. Ribosome hibernation facilitates tolerance of stationary-phase bacteria to aminoglycosides. Antimicrob Agents Chemother doi:10.1128/AAC.01532-15.

21. Wada A YY, Fujita N, Ishihama A. 1990. Structure and probable genetic location of a “ribosome modulation factor” associated with 100S ribosomes in sta tionary-phase Escherichia coli cells. Proc Natl Acad Sci doi:10.1073/pnas.87.7.2657.

22. Akiyama T WK, Schaefer R, Pratt S, Chang CB, Franklin MJ. 2017. Resuscitation of Pseudomonas aeruginosa from dormancy requires hibernation promoting factor (PA4463) for ribosome preservation. PNAS 114:3204–209.

23. Prossliner T WK, Sørensen MA, Gerdes K. 2018. Ribosome Hibernation. Annu Rev Genet doi:10.1146/annurev-genet-120215-035130.

24. (ESCMID) ESoCMaID. 2022. European Committee for Antimicrobial Susceptibility Testing (EUCAST) of the European Society of Clinical Microbiology and Infectious Dieases (ESCMID). Reading guide for broth microdilution. https://www.eucast.org/fileadmin/src/media/PDFs/EUCAST_files/Disk_test_documents/2022_manuals/Reading_guide_BMD_v_4.0_2022.pdf. Accessed

25. Trampari E HE, Wickham GJ, Ravi A, de Oliveira Martins L, Savva GM & Webber MA. 2021. Exposure of Salmonella biofilms to antibiotic concentrations rapidly selects resistance with collateral tradeoffs. . npj Biofilms Microbiomes 10.1038/s41522-020-00178-0.

26. Harrison F MA, Higgins S, Diggle SP. 2014. Development of an ex vivo porcine lung model for studying growth, virulence, and signaling of Pseudomonas aeruginosa. Infect Immun doi:10.1128/IAI.01554-14.

27. Harrington NE SE, Harrison F. 2020. Building a better biofilm - Formation of in vivo-like biofilm structures by Pseudomonas aeruginosa in a porcine model of cystic fibrosis lung infection. Biofilm 2.

28. Werthén M HL, Jensen PØ, Sternberg C, Givskov M, Bjarnsholt T. 2010. An in vitro model of bacterial infections in wounds and other soft tissues. APMIS doi:10.1111/j.1600-0463.2009.02580.x.

29. Bjarnsholt T JP, Fiandaca MJ, Pedersen J, Hansen CR, Andersen CB, Pressler T, Givskov M, Høiby N. 2009. Pseudomonas aeruginosa biofilms in the respiratory tract of cystic fibrosis patients. Pediatr Pulmonol doi:10.1002/ppul.21011.

30. Henderson AG EC, Button B, Abdullah LH, Cai LH, Leigh MW, DeMaria GC, Matsui H, Donaldson SH, Davis CW, Sheehan JK, Boucher RC, Kesimer M. 2014. Cystic fibrosis airway secretions exhibit mucin hyperconcentration and increased osmotic pressure. J Clin Invest doi:10.1172/JCI73469.

31. Baltimore RS CC, Smith GJ. 1989. Immunohistopathologic localization of Pseudomonas aeruginosa in lungs from patients with cystic fibrosis. Implications for the pathogenesis of progressive lung deterioration. Am Rev Respir Dis doi:10.1164/ajrccm/140.6.1650.

32. Reales-Calderón JA Sun Z, Mascaraque V, Pérez-Navarro E, Vialás V, Deutsch EW, Moritz RL, Gil C, Martínez JL, Molero G. 2021. A wide-ranging Pseudomonas aeruginosa PeptideAtlas build: a useful proteomic resource for a versatile pathogen. Journal of Proteomics 239.

33. Dorsey-Oresto A LT, Mosel M, Wang X, Salz T, Drlica K, Zhao X. 2013. YihE kinase is a central regulator of programmed cell death in bacteria. Cell Rep doi:10.1016/j.celrep.2013.01.026.

34. Liu Y SW, Ma L, Xu R, Yang C, Xu P, Ma C, Gao C. 2022. Metabolic Mechanism and Physiological Role of Glycerol 3-Phosphate in Pseudomonas aeruginosa PAO1. mBio doi:10.1128/mbio.02624-22.

35. Jumper J ER, Pritzel A, Green T, Figurnov M, Ronneberger O, Tunyasuvunakool K, Bates R, Žídek A, Potapenko A, Bridgland A, Meyer C, Kohl SAA, Ballard AJ, Cowie A, Romera-Paredes B, Nikolov S, Jain R, Adler J, Back T, Petersen S, Reiman D, Clancy E, Zielinski M, Steinegger M, Pacholska M, Berghammer T, Bodenstein S, Silver D, Vinyals O, Senior AW, Kavukcuoglu K, Kohli P, Hassabis D. 2021. Highly accurate protein structure prediction with AlphaFold. Nature doi:10.1038/s41586-021-03819-2.

36. Schweizer HP JR, Po C. 1997. Structure and gene-polypeptide relationships of the region encoding glycerol diffusion facilitator (glpF) and glycerol kinase (glpk) of Pseudomonas aeruginosa. Microbiology 10.1099/00221287-143-4-1287.

37. Zheng J HC, Singh VK, Martin NL, Jia Z. 2007. Crystal structure of a novel prokaryotic Ser/Thr kinase and its implication in the Cpx stress response pathway. Mol Microbiol doi:10.1111/j.1365-2958.2007.05611.x.

38. Tkachenko AG KN, Karavaeva EA, Shumkov MS. 2015. Putrescine controls the for mation of Escherichia coli persister cells tolerant to aminoglycoside netilmicin. FEMS Microbiol Lett doi:10.1111/1574-6968.12613.

39. Heijerman H WE, Conway S, Touw D, Döring G. 2009. Consensus Working Group: Inhaled medication and inhalation devices for lung diseases in patients with cystic fibrosis: a European consensus. J Cyst Fibros doi:10.1016/j.jcf.2009.04.005.

40. Harrington NE AF, Garcia-Maset R, Harrison F. 2022. Pseudomonas aeruginosa transcriptome analysis in a cystic fibrosis lung model reveals metabolic changes accompanying biofilm maturation and increased antibiotic tolerance over time. bioRxiv doi:10.1101/2022.06.30.498312:2022.06.30.498312.

41. Bielecki P PJ, Wos-Oxley ML, Loessner H, Glik J, Kawecki M, Nowak M, Tümmler B, Weiss S, dos Santos VA. 2011. In-vivo expression profiling of Pseudomonas aeruginosa infections reveals niche-specific and strain-independent transcriptional programs. PLoS One doi:10.1371/journal.pone.0024235.

42. Mavrodi D. BRF, Delaney S.M., Soule M.J., Phillips G., Thomashow L.S. 2001. Functional analysis of genes for biosynthesis of pyocyanin and phenazine-1-carboxamide from Pseudomonas aeruginosa PAO1. J Bacteriol doi:10.1128/JB.183.21.6454-6465.2001.

43. Dietrich LE TT, Price-Whelan A, Newman DK. 2008. Redox-active antibiotics control gene expression and community behavior in divergent bacteria. Science doi:10.1126/science.1160619.

44. Dietrich LE OC, Price-Whelan A, Sakhtah H, Hunter RC, Newman DK. 2013. Bacterial community morphogenesis is intimately linked to the intracellular redox state. J Bacteriol doi:10.1128/JB.02273-12.

45. Schiessl KT HF, Jo J, Nazia SZ, Wang B, Price-Whelan A, Min W, Dietrich LEP. 2019. Phenazine production promotes antibiotic tolerance and metabolic heterogeneity in Pseudomonas aeruginosa biofilms. Nat Commun doi:10.1038/s41467-019-08733-w.

46. Lin Y-C SM, Cornell WC, Silva GM, Okegbe C, Price-Whelan A, Vogel C, Dietrich LEP. 2018. Phenazines regulate Nap dependent denitrification in Pseudomonas aeruginosa biofilms. J Bacteriol 10.1128/JB.00031-18.

47. Wang Y WJ, Danhorn T, Ramos I, Croal L, Newman DK 2011. Phenazine-1-carboxylic acid promotes bacterial biofilm development via ferrous iron acquisition. J Bacteriol doi:10.1128/JB.00396-11.

48. Das T IA, Klare W, Manefield M. 2016. Role of Pyocyanin and Extracellular DNA in Facilitating Pseudomonas aeruginosa Biofilm Formation, Microbial Biofilms - Importance and Applications doi:10.5772/63497.

49. Raivio TL LS, Price NL. 2013. The Escherichia coli Cpx envelope stress response regulates genes of diverse function that impact antibiotic resistance and membrane integrity. J Bacteriol doi:10.1128/JB.00105-13.

50. Son MS MW, Kang Y, Nguyen DT, Hoang TT. 2007. In vivo evidence of Pseudomonas aeruginosa nutrient acquisition and pathogenesis in the lungs of cystic fibrosis patients. Infect Immun 10.1128/IAI.01807-06.

51. Stokes JM LA, Lobritz MA, Collins JJ. 2019. Bacterial metabolism and antibiotic efficacy. Cell Metab 10.1016/j.cmet.2019.06.009.

52. Lopatkin AJ BS, Manson AL, Stokes JM, Kohanski MA, Badran AH, Earl AM, Cheney NJ, Yang JH, Collins JJ. 2021. Clinically relevant mutations in core metabolic genes confer antibiotic resistance. Science 10.1126/science.aba0862.

53. JH M. 1972. Experiments in molecular genetics. Cold Spring Harbor Laboratory.

54. Stuart D. Woodcock KS, Richard H. Little, Danny Ward, Despoina Sifouna, James K. M. Brown, Stephen Bornemann, Jacob G. Malone. 2021. Trehalose and α-glucan mediate distinct abiotic stress responses in Pseudomonas aeruginosa. PLoS Genet 10.1371/journal.pgen.1009524.

55. M Martí BF, A Serrano-Aroca. 2018. Antimicrobial Characterization of Advanced Materials for Bioengineering Applications. J Vis Exp doi:10.3791/57710

56. Trampari E ZC, Gotts K, Savva GM, Bavro VN, Webber MA. 2022. Cefotaxime Exposure Selects Mutations within the CA-Domain of envZ Which Promote Antibiotic Resistance but Repress Biofilm Formation in Salmonella. Microbiol Spectr doi:10.1128/spectrum.02145-21.

57. Harrington NE SE, Alav I, Allen F, Moa, J, Harrison F. 2021. Antibiotic Efficacy Testing in an Ex vivo Model of Pseudomonas aeruginosa and Staphylococcus aureus Biofilms in the Cystic Fibrosis Lung. J Vis Exp doi:10.3791/62187.

58. Perez-Riverol Y BJ, Bandla C, Hewapathirana S, García-Seisdedos D, Kamatchinathan S, Kundu D, Prakash A, Frericks-Zipper A, Eisenacher M, Walzer M, Wang S, Brazma A, Vizcaíno JA. 2022. The PRIDE database resources in 2022: A Hub for mass spectrometry-based proteomics evidences. Nucleic Acids doi:10.1093/nar/gkab1038.

59. Pacheco-Moreno A SF, Ford JJ, Trippel C, Uszkoreit S, Ferrafiat L, Grenga L, Dickens R, Kelly N, Kingdon AD, Ambrosetti L, Nepogodiev SA, Findlay KC, Cheema J, Trick M, Chandra G, Tomalin G, Malone JG, Truman AW. 2021. Pan-genome analysis identifies intersecting roles for Pseudomonas specialized metabolites in potato pathogen inhibition. Elife doi:10.7554/eLife.71900.

60. Catriona M. A. Thompson JPJH, Govind Chandra, Carlo Martins, Gerhard Saalbach, Supakan Panturat, Susannah M. Bird, Samuel Ford, Richard H. Little, Ainelen Piazza, Ellie Harrison, Robert W. Jackson, Michael A. Brockhurst, Jacob G. Malone. 2023. Plasmids manipulate bacterial behaviour through translational regulatory crosstalk. PLoS Biol 10.1371/journal.pbio.3001988.

## References

1. Holloway, B.W., Genetic recombination in Pseudomonas aeruginosa. Journal of General Microbiology, 1955. 13(3): p. 572–81.

2. Woodcock, D.M., et al., Quantitative evaluation of Escherichia coli host strains for tolerance to cytosine methylation in plasmid and phage recombinants. Nucleic Acids Res, 1989. 17(9): p. 3469–78.

3. Karimova G, Pidoux J, Ullmann A, Ladant D. 1998. A bacterial two-hybrid system based on a reconstituted signal transduction pathway. Proc Natl Acad Sci U S A 95:5752–5756.

4. Scott, T.A., et al., An L-threonine transaldolase is required for L-threo-beta-hydroxy-alpha-amino acid assembly during obafluorin biosynthesis. Nature Communications, 2017. 8.

5. Heeb, S., et al., Small, stable shuttle vectors based on the minimal pVS1 replicon for use in gram-negative, plant-associated bacteria. Mol Plant Microbe Interact, 2000. 13(2): p. 232–7.

